# Loss of cardiomyocyte eukaryotic elongation factor 1A2 in mice triggers cardiomyopathy due to defective proteostasis

**DOI:** 10.64898/2026.01.14.699518

**Authors:** Abel Martin-Garrido, Nadine Weyrauch, Jorge Ruiz-Orera, Jeanette Eresch, Sonja Reitter, Julio Cordero, Frauke Senger, Viktoria Scheich, Andrea Grund, Merve Keles, Nina Weinzierl, Ellen Hofmann, Felix A. Trogisch, Shruthi Hemanna, Michael Buettner, Gernot Poschet, Mandy Rettel, Frank Stein, Eva A. Rog-Zielinska, Norbert Frey, Georg Stoecklin, Mirko Völkers, Norbert Hübner, Gergana Dobreva, Joerg Heineke

## Abstract

Eukaryotic elongation factor 1A (eEF1A) delivers aminoacyl-tRNAs to ribosomes but also has additional, non-canonical functions. Mammals express two paralogs: eEF1A1 is ubiquitous, whereas eEF1A2 is confined to adult cardiomyocytes, skeletal myocytes, and neurons. Mutations in EEF1A2 cause cardiomyopathy, but underlying mechanisms remain unclear. Using adult, cardiomyocyte-specific *Eef1a2* knock-out (Eef1a2-cKO) and *Eef1a1*/*Eef1a2* double knock-out mice, we show that Eef1a2-cKO animals develop cardiomyopathy with increased mortality, systolic dysfunction, and fibrosis, despite unchanged global protein synthesis, while double knock-out mice die early in a sudden manner. Multi-omics analyses reveal post-transcriptional upregulation of ribosomal proteins and translational regulators in both models. Eef1a2-cKO hearts accumulate autophagosomes and protein aggregates, indicating defective autophagy. Mechanistically, we found that eEF1A2 functions as a chaperone supporting protein folding and proteostasis in cardiomyocytes. Early Rapamycin treatment (mTORC1 inhibition) normalizes systolic heart function and survival in Eef1a2-cKO mice and clears autophagosomes and protein aggregates. Thus, eEF1A2 maintains cardiac proteostasis, and mTORC1 inhibition may represent a therapeutic strategy for patients with EEF1A2 mutations.

## Introduction

The eukaryotic translation elongation factor 1A (eEF1A) is expressed as two paralogous isoforms, eEF1A1 and eEF1A2, which share a high degree of sequence homology (92% at the mRNA level and 98% at the protein level) (1). Its canonical role is to facilitate the GTP-dependent delivery of aminoacyl– tRNAs to ribosomes, a process that is essential for the fidelity and efficiency of polypeptide synthesis during the elongation phase of translation (2, 3).

During postnatal development, a well-characterized isoform switch occurs in skeletal muscle cells, neurons, and cardiomyocytes, where fetally expressed eEF1A1 is replaced by the adult paralog eEF1A2 (4–6). In contrast, most other cell types continue to express eEF1A1 throughout adulthood. The functional importance of this developmental switch was first demonstrated by an elegant study of Cathy Abbott’s lab, which showed that the “wasted” mutation –an autosomal recessive deletion of the *Eef1a2* gene– results in weight loss, progressive paralysis, and premature death around 28 days after birth (5). More recently, Feng et al. provided further evidence for the essential role of eEF1A2 by demonstrating that cardiomyocyte-specific neonatal deletion of *Eef1a2* leads to cardiomyopathy and premature death between 7 and 22 weeks after birth in mice (7).

eEF1A is one of the most abundant intracellular proteins, accounting for approximately 1 to 11% of the total cellular proteome (3). A previous study in rabbit reticulocytes has estimated an eEF1A-to-ribosome ratio of approximately 25:1 and an eEF1A-to-tRNA ratio of 13:1, indicating that free eEF1A is the most prevalent form (3). Given its abundance, it is not surprising that eEF1A is also involved in several other cellular processes, including actin polymerization, proteasome activity, viral replication, protein folding, and cell proliferation (8). Notably, eEF1A2 exhibits translation buffering, in which protein levels remain stable despite changes in mRNA expression. This phenomenon is observed in human and non-human primateś hearts, highlighting the importance of maintaining its protein expression (9).

In 2017, Cao et al. reported the first known case of a homozygous *EEF1A2* missense mutation (P333L) in humans, associated with cardiomyopathy, epilepsy, global developmental delay, and early mortality (10). In addition, eEF1A was implicated in the pathogenesis of cardiac hypertrophy (11). Feng et al. generated a knock-in mouse model, in which the endogenous cardiac *Eef1a2* gene was replaced with the P333L variant (7). Despite exhibiting *Eef1a2* mRNA levels comparable to wild-type (WT) controls, mutant mice showed a marked reduction in eEF1A2 protein levels in cardiac tissue, suggesting that the P333L substitution acts as a loss-of-function mutation that compromises protein stability and activity. Indeed, cardiac-specific P333L-*Eef1a2* mice and neonatal onset *Eef1a2* knockout (KO) mice were phenotypically indistinguishable. Both lines began to exhibit mortality from 8 weeks of age, with complete lethality by 17 weeks. Interestingly, global protein synthesis was unaffected in P333L-*Eef1a2* and *Eef1a2* KO hearts, suggesting that the cardiomyopathy observed in both models might be independent of the canonical function of eEF1A2. Thus, the molecular mechanisms by which loss of eEF1A2 leads to heart disease remains unknown.

In this study, we investigate the role of eEF1A in adult cardiomyocytes. Our results show that cardiomyocyte-specific deletion of *Eef1a2* in adult mice leads to cardiomyopathy, with a lethality rate of 45% two months after gene deletion. Furthermore, we demonstrate that eEF1A1 plays a protective role in the absence of eEF1A2, as mice with cardiomyocyte-specific deletion of both *Eef1a1* and *Eef1a2* exhibit complete lethality within six weeks of gene deletion. Mechanistically, by combining mRNA sequencing, proteomics, and ribosomal sequencing analyses, we demonstrate that in eEF1A2-deficient hearts, proteins encoded by mRNAs with a 5′-terminal oligopyrimidine (TOP) motif are specifically upregulated. In addition, we identify eEF1A2 as a critical chaperone involved in cardiac protein folding. Interestingly, treatment with Rapamycin rescued cardiac dysfunction and reduced the accumulation of misfolded proteins caused by *Eef1a2* deletion, opening an avenue for a tailored therapeutic approach for patients with inherited *EEF1A2* mutations.

## Results

### Induced deletion of *Eef1a2* in adult cardiomyocytes leads to cardiomyopathy

Previous findings indicate that *Eef1a2* expression is essential for maintaining cardiac function during postnatal heart development. However, the role of eEF1A2 in adult cardiomyocytes remains unknown (7). To investigate the function of eEF1A2 in the adult heart, we generated a cardiomyocyte-specific, Tamoxifen-inducible conditional knockout mouse model (Eef1a2-cKO) by interbreeding αMHC-MerCreMer with newly generated Eef1a2 flox/flox mice. Tamoxifen was administered to mice aged 6– 7 weeks to induce Cre recombinase activity and specifically delete *Eef1a2* in cardiomyocytes. Eef1a2 flox/flox mice (littermates) were used as control (Eef1a2-WT). Our results show that Eef1a2-cKO mice began to exhibit mortality at 6 weeks after Tamoxifen injection, with a 55% survival rate observed at two months (Figure 1A). Cardiac systolic function, assessed as ejection fraction in echocardiography, showed no significant differences between control and Eef1a2-cKO mice at one month after Tamoxifen injections, but by two months, Eef1a2-cKO hearts displayed marked cardiac dysfunction, with an approximately 60% reduction in left ventricular ejection fraction compared to control mice (Figure 1B, Figure S1A, Table 1). Gravimetric analysis revealed significant enlargement of both ventricles and atria in Eef1a2-cKO mice (Figure 1C), which were not detectable one month earlier (Figure S1B). Cardiac hypertrophy in Eef1a2-cKO mice was accompanied by an increase in cardiomyocyte cross-sectional area, indicative of cellular hypertrophy (Figure 1D). Furthermore, Sirius red staining showed advanced myocardial fibrosis two months after Tamoxifen injections in Eef1a2-cKO mice, which started to emerge already at the one month time point (Figure 1E, S1D). Analysis of hypertrophic gene expression revealed a significant dysregulation as early as one month after Tamoxifen administration (Figure S1E), which was further enhanced one month later (Figure 1F). Male and female Eef1a2-cKO mice displayed similar changes in echocardiography and morphometry (Table S1). Finally, given that eEF1A2 is canonically involved in the elongation phase of protein synthesis, we investigated whether its deletion affects global protein synthesis. Using a puromycin incorporation assay, we found no significant differences in global protein synthesis between isolated cardiomyocytes from Eef1a2-cKO and control mice at one or two months post gene deletion (Figure 1G-H and S1F). Viability, cardiac function, and morphology were similar to WT mice with only minor differences 4 months after Tamoxifen administration in αMHC-MerCreMer mice on a WT background, demonstrating that the effects observed in Eef1a2-cKO mice were due to the deletion of *Eef1a2*, and not to Tamoxifen-induced Cre activation (Tables S2-S3). Collectively, our results show that deletion of eEF1A2 in adult cardiomyocytes induces progressive cardiac dysfunction and mouse mortality between one and two months after initiating deletion of the gene, while general protein synthesis activity is not affected.

**Figure 1.**
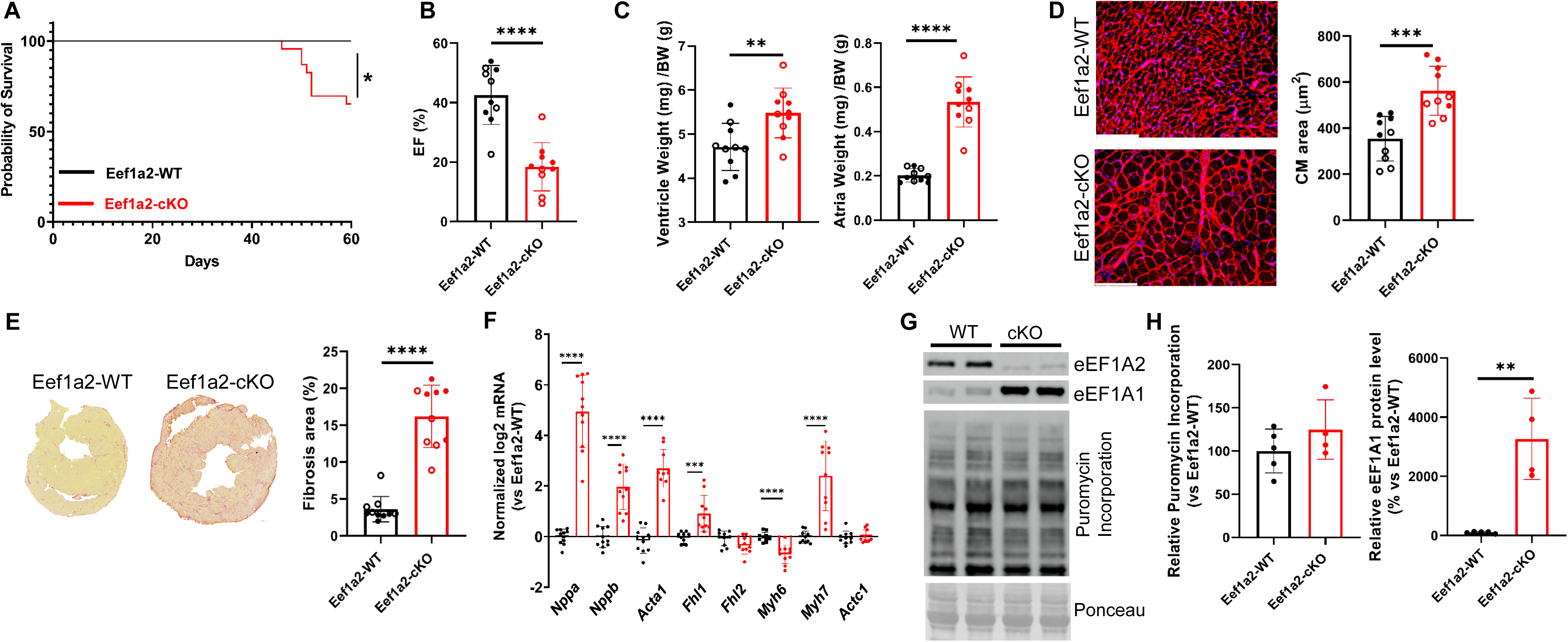
Effects of cardiomyocyte-specific Eef1a2 deletion. **(A)** Kaplan–Meier survival curves of Eef1a2-WT (n = 10, 10/10 survived) and Eef1a2-cKO (n = 16, 10/16 survived) mice. *P < 0.05, log-rank (Mantel–Cox) test. (**B**) Left ventricular ejection fraction (EF) of Eef1a2-WT (n = 10) and Eef1a2-cKO (n = 10) mice. ****P < 0.0001, unpaired two-tailed t-test. (**C**) Ventricular (left) and atrial (right) weight normalized to body weight in Eef1a2-WT (n = 10) and Eef1a2-cKO (n = 10) mice. **P < 0.01, ****P < 0.0001; unpaired two-tailed t-test. (**D**) Representative microscopic images of cardiac sections stained with wheat germ agglutinin (WGA) and corresponding quantification of cardiomyocyte (CM) cross-sectional area from Eef1a2-WT (n = 10) and Eef1a2-cKO (n = 10) mice (scale bars, 100 µm). ***P < 0.001, unpaired two-tailed t-test. (**E**) Representative Sirius Red staining of cardiac fibrosis and corresponding quantification of fibrotic area from Eef1a2-WT (n = 10) and Eef1a2-cKO (n = 10) mice. ****P < 0.0001, Mann–Whitney-U-test. (**F)** Normalized mRNA expression levels of selected genes in cardiac tissue from Eef1a2-WT (n = 10) and Eef1a2-cKO (n = 10) mice. ***P < 0.001, ****P < 0.0001; unpaired two-tailed t-test. (**G**) Representative Western blot analysis of isolated cardiomyocytes. (**H**) Quantification of puromycin incorporation and eEF1A1 protein levels in isolated cardiomyocytes from Eef1a2-WT (n = 5) and Eef1a2-cKO (n = 4) mice from independent isolations. **P < 0.01; unpaired two-tailed t-test. Data in B-H were obtained 2 months after Tamoxifen administration. Filled circles denote males; open circles denote females.

**Table 1:**
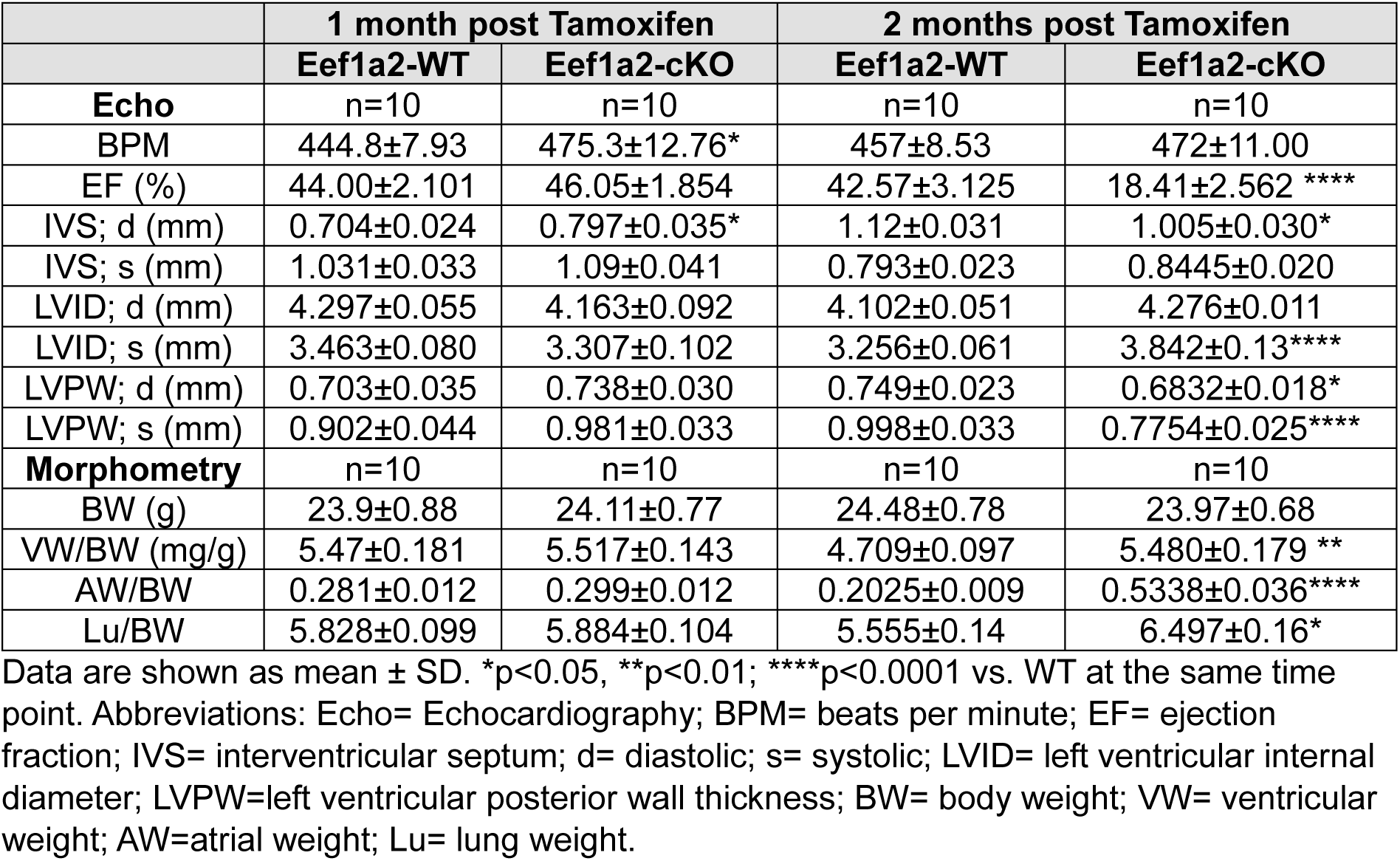
Echocardiography and morphometry in Eef1a2-WT and Eef1a2-cKO mice.

|  | 1 month post Tamoxifen |  | 2 months post Tamoxifen |  |
| --- | --- | --- | --- | --- |
|  | Eef1a2-WT | Eef1a2-cKO | Eef1a2-WT | Eef1a2-cKO |
| **Echo** | n=10 | n=10 | n=10 | n=10 |
| BPM | 444.8±7.93 | 475.3±12.76* | 457±8.53 | 472±11.00 |
| EF (%) | 44.00±2.101 | 46.05±1.854 | 42.57±3.125 | 18.41±2.562 **** |
| IVS; d (mm) | 0.704±0.024 | 0.797±0.035* | 1.12±0.031 | 1.005±0.030* |
| IVS; s (mm) | 1.031±0.033 | 1.09±0.041 | 0.793±0.023 | 0.8445±0.020 |
| LVID; d (mm) | 4.297±0.055 | 4.163±0.092 | 4.102±0.051 | 4.276±0.011 |
| LVID; s (mm) | 3.463±0.080 | 3.307±0.102 | 3.256±0.061 | 3.842±0.13**** |
| LVPW; d (mm) | 0.703±0.035 | 0.738±0.030 | 0.749±0.023 | 0.6832±0.018* |
| LVPW; s (mm) | 0.902±0.044 | 0.981±0.033 | 0.998±0.033 | 0.7754±0.025**** |
| **Morphometry** | n=10 | n=10 | n=10 | n=10 |
| BW (g) | 23.9±0.88 | 24.11±0.77 | 24.48±0.78 | 23.97±0.68 |
| VW/BW (mg/g) | 5.47±0.181 | 5.517±0.143 | 4.709±0.097 | 5.480±0.179 ** |
| AW/BW | 0.281±0.012 | 0.299±0.012 | 0.2025±0.009 | 0.5338±0.036**** |
| Lu/BW | 5.828±0.099 | 5.884±0.104 | 5.555±0.14 | 6.497±0.16* |

Data are shown as mean ± SD. \*p<0.05, \*\*p<0.01; \*\*\*\*p<0.0001 vs. WT at the same time point. Abbreviations: Echo= Echocardiography; BPM= beats per minute; EF= ejection fraction; IVS= interventricular septum; d= diastolic; s= systolic; LVID= left ventricular internal diameter; LVPW=left ventricular posterior wall thickness; BW= body weight; VW= ventricular weight; AW=atrial weight; Lu= lung weight.

### Combined adult-onset ablation of *Eef1a1* and *Eef1a2* leads to early sudden death

Western blot analysis of the different eEF1A paralogs in the heart revealed that a compensatory upregulation of eEF1A1 was detectable in isolated cardiomyocytes upon *Eef1a2* deletion as early as one month after Tamoxifen administration (Figure 1G-H and S1F). Interestingly, this increase appeared to be regulated post-transcriptionally, as mRNA levels of *Eef1a1* remained similar between isolated cardiomyocytes from WT and Eef1a2-KO mice (Figure S1G). To assess the functional significance of this upregulation, we generated cardiomyocyte-specific, Tamoxifen-inducible conditional double knockout mice lacking both *Eef1a1* and *Eef1a2* (Eef1a1/Eef1a2-cKO). Our results show 100% mortality between 40- and 45-days post-Tamoxifen in Eef1a1/Eef1a2-cKO mice (Figure 2A). Interestingly, echocardiographic assessment at one month post-Tamoxifen, showed no signs of cardiac dysfunction, but a small reduction in diastolic and systolic left ventricular dimensions in double knockout mice, which occurred similarly in both sexes (Figure 2B and Table 2 and Table S4). In line with this observation, gravimetric analysis (Figure 2C) and morphological assessment of cardiomyocyte cross-sectional area (Figure 2D) revealed no significant differences between Eef1a1/Eef1a2-cKO and control mice. Furthermore, no cardiac fibrosis was observed in the double knock-out hearts at one month post-Tamoxifen, around 10 days before mortality started (Figure 2E). However, analysis of hypertrophic gene expression revealed a significant dysregulation of all analyzed genes (Figure 2F), which appeared to be more pronounced than that observed in Eef1a2-cKO mice at the one month time point (Figure S1E). Notably, in contrast to the single *Eef1a2* knockout, puromycin incorporation was significantly reduced by about 30% in cardiomyocytes from Eef1a1/Eef1a2-cKO mice, indicating mildly impaired global protein synthesis (Figure 2G-H).

**Figure 2.**
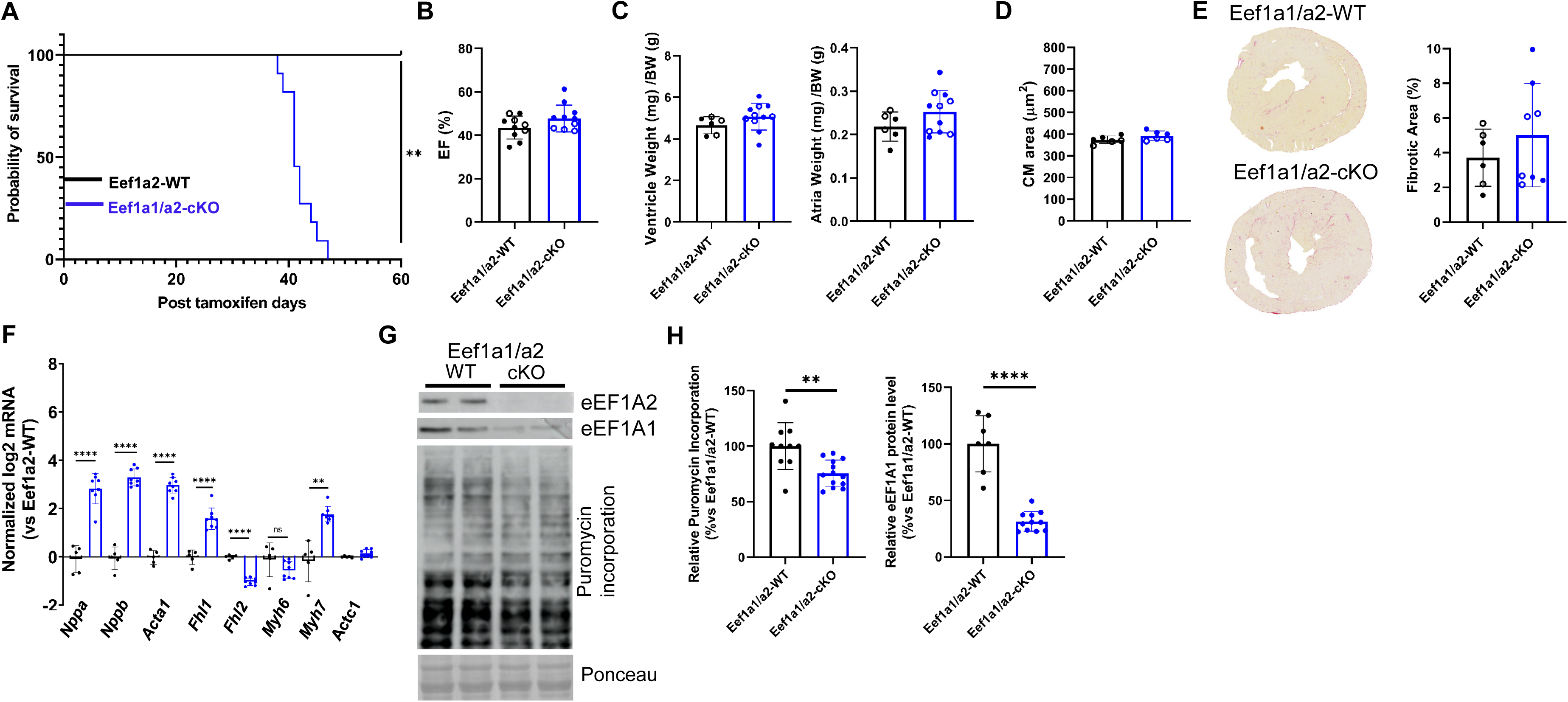
Effects of cardiomyocyte-specific Eef1a1 and Eef1a2 deletion. **(A)** Kaplan–Meier survival curves of Eef1a1/a2-WT (n = 5; 5/5 survived) and Eef1a1/a2-cKO (n = 11; 0/11 survived) mice. **P < 0.01, log-rank (Mantel–Cox) test. (**B**) Left ventricular ejection fraction (EF) of Eef1a1/a2-WT (n = 10) and Eef1a1/a2-cKO (n = 11). (**C**) Ventricular (left) and atrial (right) weight normalized to body weight in Eef1a1/a2-WT (n = 6) and Eef1a1/a2-cKO (n = 11) mice at 1 month after Eef1a1/a2 deletion. (**D**) Quantification of cardiomyocyte (CM) cross-sectional area in Eef1a1/a2-WT (n = 6) and Eef1a1/a2-cKO (n = 6) mice. (**E**) Representative Sirius Red staining of cardiac fibrosis and corresponding quantification of fibrotic area in Eef1a1/a2-WT (n = 6) and Eef1a1/a2-cKO (n = 8) mice (**F**) Normalized mRNA expression levels of selected genes in cardiac tissue from Eef1a1/a2-WT (n = 5) and Eef1a1/a2-cKO (n = 8) mice at 1 month after Eef1a1/a2 deletion. (**G**) Representative Western blot analysis of isolated cardiomyocytes. Ponceau staining shows equal protein loading and blotting. (**H**) Quantification of eEF1A1 expression and puromycin incorporation in isolated cardiomyocytes from Eef1a1/a2-WT (Eef1a1, n = 7 independent isolations; puromycin, n = 10 independent isolations) and Eef1a1/a2-cKO mice (Eef1a1, n = 6; puromycin, n = 11). ****P < 0.0001, **P < 0.01; unpaired two-tailed t-test. Data in B-H were obtained one month after Tamoxifen administration. Filled circles denote males; open circles denote females.

**Table 2:** Echocardiography and morphometry in Eef1a1/a2-WT and Eef1a1/a2-cKO mice.

|  | Eef1a1/a2-WT | Eef1a1/a2-cKO |
| --- | --- | --- |
| **Echo** | n=10 | n=11 |
| BPM | 450±5.71 | 442.8±6.04 |
| EF (%) | 43.60±1.67 | 47.75±1.86 |
| IVS;d (mm) | 0.7837±0.021 | 0.7666±0.027 |
| IVS;s (mm) | 1.091±0.035 | 1.135±0.034 |
| LVID;d (mm) | 4.454±0.07 | 4.15±0.061** |
| LVID;s (mm) | 3.588±0.07 | 3.217±0.049*** |
| LVPW;d (mm) | 0.8283±0.033 | 0.7454±0.029 |
| LVPW;s (mm) | 1.049±0.057 | 1.026±0.053 |
| **Morphometry** | n=6 | n=11 |
| BW (g) | 21.81±1.18 | 21.88±0.78 |
| VW/BW (mg/g) | 4.658±0.16 | 5.07±0.19 |
| AW/BW | 0.219±0.013 | 0.2524±0.014 |
| Lu/BW | 6.325±0.34 | 6.947±0.30 |

Animals are analyzed one month after Tamoxifen administration. Data are shown as mean ± SD. \*\*p<0.01; \*\*\*p<0.001 vs. WT at the same time point.

We then analyzed mice with adult induced cardiomyocyte-specific *Eef1a1* knock-out (Eef1a1-cKO) and found that viability, cardiac function, and morphology remained unchanged in mice of both sexes at four months after Tamoxifen application compared to control mice (Figure S2, Tables S5-6). Moreover, puromycin incorporation and eEF1A2 expression were similar between control and Eef1a1-cKO isolated cardiomyocytes one month after Tamoxifen administration (Figure S2E). These results clearly demonstrate that cardiac eEF1A1 expression is dispensable under physiological conditions in adulthood, when eEF1A2 is the predominant paralog. Furthermore, our data suggest that the upregulation of eEF1A1 observed in Eef1a2-cKO mice is a compensatory mechanism, and that eEF1A1 and eEF1A2 can at least partially compensate for each other in cardiomyocytes, particularly in maintaining protein synthesis.

### Increased translation of ribosomal proteins upon *Eef1a2* gene deletion

Next, we investigated potential mechanisms underlying the cardiac dysfunction induced by *Eef1a2* knock-out. Proteomic and RNA-seq analyses were performed on whole heart tissue one month after Tamoxifen mediated gene deletion, in order to capture initial molecular changes that occur prior to the onset of cardiac dysfunction. RNA-seq analysis revealed upregulation of muscle, sarcomeric, hypertrophic and unfolded protein response genes as well as downregulation of genes involved in action potential generation, endothelial proliferation, muscle contraction and potassium transport (Figure S3A-B). Interestingly, proteomic analysis uncovered a marked upregulation of ribosomal proteins and translation regulatory factors, as well as a downregulation of proteins related to lipid oxidation, fatty acid and carboxylic acid metabolic processes (Figure S3C-D).

Given the heterogeneous cell composition of the heart, we next sought to validate these proteomic findings in isolated cardiomyocytes. For this purpose, we isolated adult cardiomyocytes from Eef1a2-cKO, Eef1a1/a2-cKO and the respective control mice one month after Tamoxifen injections and subjected them to proteomic analyses (Figure 3A). Proteins identified in cardiomyocytes of both knock-out mouse lines were also compared with the results from cardiomyocytes isolated from αMHC-MerCreMer transgenic mice (Figure S4A), and those expressed at similar levels in all groups were removed from the list. As shown in Figure 3B, we identified 40 differentially expressed proteins in Eef1a2-cKO, and 301 in Eef1a1/Eef1a2 double-cKO cardiomyocytes (Figure 3D). Gene Ontology (GO) analysis confirmed an upregulation of ribosomal proteins and translation regulatory factors in Eef1a2-cKO, which was even more pronounced in the double-cKO cardiomyocytes (Figure 3C and E, and Figure S4B-C). Notably, 95% of the differentially expressed proteins in the Eef1a2-cKO group were also observed in the double-cKO, while they represented only 20% of the total changes seen in the double knock-out (Figure 3F). Western blot analysis confirmed an upregulation of eEF2 and RPS6 proteins in isolated cardiomyocytes from Eef1a2-cKO and Eef1a1/a2-cKO mice within one month after Tamoxifen administration (Figure 3G-H). In contrast, mRNA levels of *Rps6* and *Eef2* were not significantly changed (Figure S4D).

**Figure 3.**
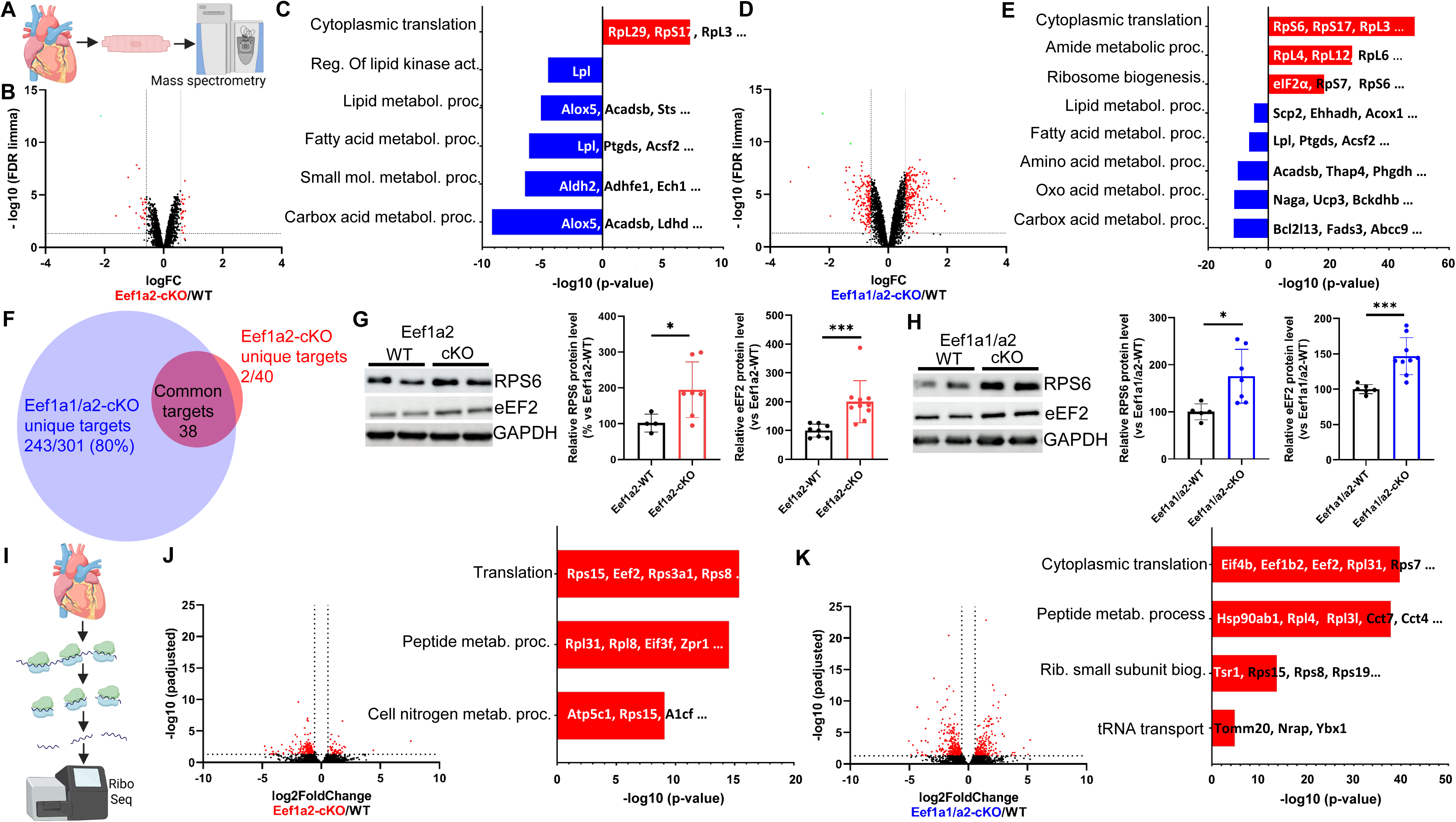
Effects of cardiomyocyte-specific Eef1a2 and Eef1a1/a2 deletion on the proteome and translation. **(A)** Schematic representation of the proteomic analysis workflow. (**B**) Volcano plot showing fold change (FC; log₂) and P value of protein abundance changes in isolated cardiomyocytes from WT (n = 4) and Eef1a2-cKO (n = 4) mice at 1 month after Eef1a2 deletion. Significantly changed proteins are highlighted in red. (**C**) Gene Ontology (GO) analysis of significant proteins identified in (B). Upregulated or downregulated GO terms in mutant mice are shown in red and blue, respectively. (**D**) Volcano plot showing fold change (FC; log₂) and P value of protein abundance changes in isolated cardiomyocytes from WT (n = 4) and Eef1a1/a2-cKO (n = 4) mice at 1 month after Eef1a1/a2 deletion. Significantly changed proteins are highlighted in red. (**E**) GO analysis of significant proteins identified in (D). Upregulated or downregulated GO terms in mutant mice are shown in red and blue, respectively. (**F**) Venn diagram showing overlap of significant proteins identified in (B) and (D). (**G**) Representative Western blot analysis and quantification of RPS6 and eEF2 in isolated cardiomyocytes from Eef1a2-WT (RPS6, n = 4; eEF2, n = 8) and Eef1a2-cKO (RPS6, n = 7; eEF2, n = 10) mice. *P < 0.05, ***P < 0.001; unpaired two-tailed t-test. (**H**) Representative Western blot analysis and quantification of indicated proteins in isolated cardiomyocytes from Eef1a1/a2-WT (RPS6, n = 5; eEF2, n = 6) and Eef1a1/a2-cKO (RPS6, n = 7; eEF2, n = 9) mice. *P < 0.05, ***P < 0.001; unpaired two-tailed t-test. (**I**) Schematic representation of the Ribo-seq analysis workflow. (**J**) Volcano plot showing fold change and P value of changes in translation efficiency in cardiac tissue from WT (n = 4) and Eef1a2-cKO (n = 4) mice at 1 month after Eef1a2 deletion. Regulated genes are highlighted in red, with corresponding GO analysis of upregulated genes in mutant mice. (**K**) Volcano plot showing fold change and P value of changes in translation efficiency in cardiac tissue from WT (n = 4) and Eef1a1/a2-cKO (n = 4) mice at 1 month after Eef1a1/a2 deletion. Regulated genes are highlighted in red, with corresponding GO analysis shown of upregulated genes in mutant mice.

Since upregulation of eEF2 and RPS6 occurred predominantly at the protein and not the mRNA level, the effect on ribosomal proteins and translation regulatory factors appeared to be mediated mainly post-transcriptionally. To verify this, we performed ribosomal sequencing (Ribo-Seq) to measure ribosome-protected mRNA fragments in isolated hearts one month after Tamoxifen injections (Figure 3I). Ribo-Seq revealed the highest increase in ribosome protected fragments for ribosomal protein and translation factor mRNAs in Eef1a2- and Eef1a1/Eef1a2-cKO hearts compared to controls (Figure S5A-B). More importantly, when translational efficiency (TE) was calculated by normalizing ribosome protected fragments of a gene to its mRNA expression in the same sample, ribosomal proteins and translation factors were again among the most upregulated gene groups in both knock-out mouse lines (Figure 3J-K). Of note, a principal component analysis of TE values clearly distinguished each experimental group (Figure S6A). Consistent with our proteomics findings, we observed a stronger increase in TE for ribosomal proteins and translation factors in the Eef1a1/Eef1a2-cKO hearts compared with the Eef1a2-cKO hearts (Figure S5C). This dataset was also used for an analysis of mRNA codon occupancy. While no major differences were observed in Eef1a2-cKO hearts, Asparagine (N), Glycine (G) and Leucine (L) codon occupancy appeared slightly reduced in Eef1a1/Eef1a2-cKO, and Lysine (K), Proline (P) as well as Arginine (R) codon occupancy was increased. This result suggested that N, G and L codons are more efficiently decoded, whereas K, P and R codons are decoded more slowly in Eef1a1/Eef1a2-cKO hearts (Figure S6B). Collectively, these results provide evidence that eEF1A2 specifically controls the polypeptide synthesis of ribosomal proteins and translation factors in cardiomyocytes.

Next, we investigated whether the upregulation of ribosomal proteins and elongation factors altered the polysome distribution or the spatial regulation of protein synthesis, as this had been previously shown to be critical for cardiac function (12–14). We separated monosome and polysome fractions using sucrose density gradient ultracentrifugation, yet no differences were observed in the fraction of polysomes between control, Eef1a2-cKO and Eef1a1/Eef1a2-cKO hearts (Figure S7A-B). Similarly, the distribution of RPS6 (a representative ribosomal small subunit protein), eEF2 (a representative elongation factor), eIF2α (a representative initiation factor) and EDF1 (a marker of ribosome collisions (15, 16)) across monosome and polysome fractions was similar in the different genotypes (Figure S7C-D). The Z-disc has been reported to serve as an anchor for the protein synthesis machinery in adult cardiomyocytes. Accordingly, staining for eEF1A2 revealed a sarcomeric staining pattern, which was lost in Eef1a2-cKO cardiomyocytes (Figure S8A). Immunofluorescence analysis of the ribosomal protein RPS6 together with the Z-disc protein α-actinin in isolated cardiomyocytes from Eef1a1/a2-cKO mice revealed no differences in RPS6 localization compared with WT controls (Figure S8B). In summary, our results indicate that the protein synthesis machinery and its subcellular localization remain unaltered after deletion of eEF1A2, suggesting that the deleterious cardiac effects observed are not due to mislocalization of protein synthesis.

All mRNAs encoding ribosomal proteins, as well as many proteins encoding translation initiation and elongation factors, contain a TOP motif at their 5′ end, hereafter termed 5′TOP mRNAs (17). Multiple studies have shown that the translation of 5′TOP mRNAs is regulated by mTORC1 activity (18–21). We therefore investigated whether mTORC1 activity is affected by the deletion of eEF1A2, using phosphorylation of RPS6 at Ser240/244 as a commonly used surrogate marker for mTORC1 activity, given that this phosphorylation is largely dependent on the mTORC1–S6K1 signaling pathway. To explore the potential involvement of mTORC1 in cardiac dysfunction observed in Eef1a2-cKO mice, we analyzed heart samples collected two months after eEF1A2 deletion, when cardiac dysfunction was evident. Indeed, both total RPS6 as well as relative phosphorylation of RPS6 were upregulated at this time point (Figure 4A-B). Moreover, immunofluorescence staining confirmed that phosphorylation of RPS6 occurred in cardiomyocytes by co-staining phospho-RPS6 with cardiomyocyte specific α-actinin (Figure 4C). Notably, eEF2 was also upregulated in Eef1a2-cKO cardiac tissue, but the ratio of phospho-versus total eEF2 remained unaltered (Figure S9A-B), in line with the notion that eEF2 phosphorylation is not controlled by the mTORC1 pathway.

**Figure 4.**
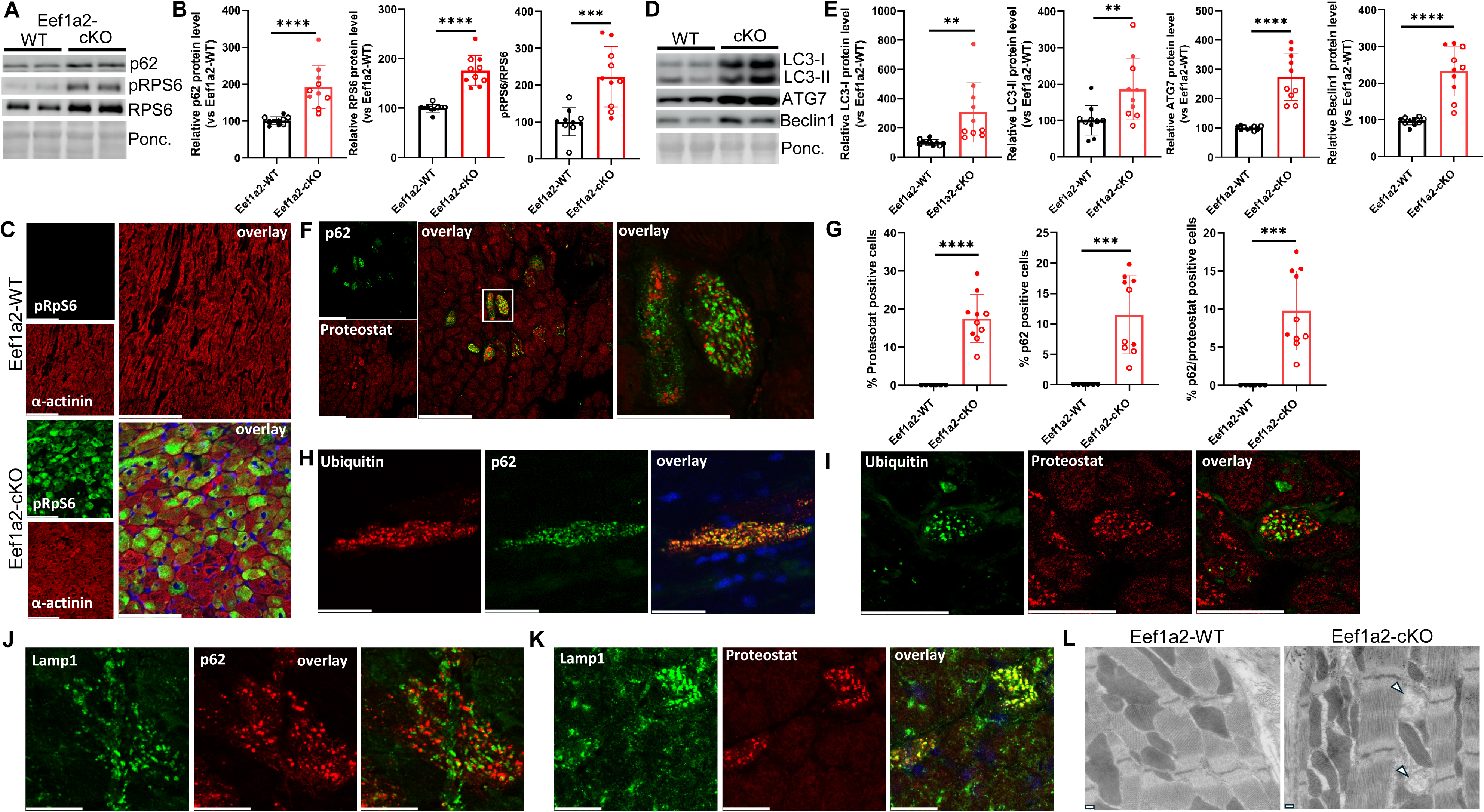
Effects of cardiomyocyte-specific Eef1a2 deletion on autophagy. (**A**) Representative Western blot analysis of the indicated proteins in cardiac tissue from Eef1a2-WT and Eef1a2-cKO mice. Ponceau (Ponc.) staining shows equal protein loading and blotting. (**B**) Quantification of the proteins shown in (A) from Eef1a2-WT (n = 9) and Eef1a2-cKO (n = 10) mice. ***P < 0.001, ****P < 0.0001; unpaired two-tailed t-test. (**C**) Representative co-immunostaining of the indicated proteins in cardiac tissue from Eef1a2-WT or Eef1a2-cKO mice (scale bars, 100 µm). (**D**) Representative Western blot analysis of indicated proteins in cardiac tissue from Eef1a2-WT and Eef1a2-cKO mice. (**E**) Quantification of the proteins shown in (D) from Eef1a2-WT (n = 9) and Eef1a2-cKO (n = 10) mice. **P < 0.01, ****P < 0.0001; unpaired two-tailed t-test. (**F**) Representative co-immunostaining of Proteostat (red) and p62 (green) in cardiac tissue from Eef1a2-cKO (left; scale bars, 100 µm). The right panel shows a magnified view of the boxed region (scale bar, 40 µm). (**G**) Quantification of cardiomyoycte immunostaining from (F) from Eef1a2-WT (n = 6) and Eef1a2-cKO (n = 10) mice. ***P < 0.001, ****P < 0.0001; unpaired two-tailed t-test. (**H**) Representative co-immunostaining of indicated proteins in cardiac tissue from Eef1a2-cKO mice (scale bars, 40 µm). (**I - K**) Representative co-immunostaining as indicated in cardiac tissue from Eef1a2-cKO mice (scale bars, 40 µm). (**L**) Representative transmission electron microscopy images of cardiac tissue from Eef1a2-WT or Eef1a2-cKO mice (scale bars, 200 nm). Arrowheads point at autophagosomes. All data were obtained 2 months after Tamoxifen administration. Filled circles denote males; open circles denote females.

Since proteomics analysis of isolated cardiomyocytes and cardiac tissue (Figures 3B-E and S3D) revealed a significant downregulation of proteins involved in carboxylic acid metabolism, we next investigated the metabolic consequences of eEF1A2 deletion. In metabolomic studies of cardiac tissue from Eef1a2-cKO mice, Gene Ontology (GO) analysis showed a specific downregulation of serine, methionine, glycine, and isoleucine metabolism (Figures S10A-B), which was also found in the myocardium of Eef1a1/a2-cKO mice, whereas no significant differences were observed for the remaining detected amino acids compared to controls (Figure S10B). These changes might relate to increased translational imitation due to increased mTORC1 activation and therefore increased amino acid (especially methionine) usage.

### Accumulation of autophagosomes and misfolded proteins in the myocardium of Eef1a2-cKO mice

Autophagy is an essential process responsible for the removal of misfolded or aggregated proteins and the clearance of damaged organelles. Previous studies have shown that mTORC1 activity negatively regulates autophagy (22). p62 is an adaptor protein involved in selective autophagy playing a key role in the clearance of protein aggregates and organelles via macroautophagy (23). As shown in Figure 4A-B, we observed an increase in p62 protein levels in Eef1a2-cKO mouse hearts, suggesting impaired macroautophagic activity (24). Macroautophagy is a multistep and complex process, wherein the conversion of cytosolic LC3B (LC3-I) to its lipidated, autophagosome-associated form (LC3-II) is a crucial step. Interestingly, the levels of LC3-I and LC3-II were higher in cardiac tissue of Eef1a2-cKO mice (Figure 4D-E) and we observed an increase in Beclin1 and Atg7 (Figure 4D-E), critical components of the multiprotein complex that participates in vesicle nucleation and the processing of LC3-I to LC3-II. Using immunofluorescence, we detected p62 positive puncta in around 10-20% of the cardiomyocytes in hearts of Eef1a2-cKO mice, but not in controls (Figure 4F-G). Given that our results suggest a deficit in macroautophagic activity in Eef1a2-cKO hearts, we next investigated whether protein aggregates accumulate. Using a specific probe (Proteostat) to detect aggregated proteins, we observed that 20– 30% of cardiomyocytes in Eef1a2-cKO hearts contained protein aggregates, whereas none were detected in control hearts (Figure 4F-G). Moreover, co-staining with p62 in cardiac tissue revealed that approximately 50–60% of Proteostat-positive cardiomyocytes were also positive for p62. Conversely, among the p62-positive cardiomyocytes (10–20% of the total cardiomyocytes), around 70% were also Proteostat-positive (Figure S9C-D). p62-and Proteostat-positive structures did not colocalize, even in cardiomyocytes positive for both markers, indicating that these signals represent distinct intracellular entities (Figure 4F). Next, we further characterized the properties of the two structures identified by p62 and Proteostat staining. To this end, we performed co-immunostaining of p62 or Proteostat with an anti-ubiquitin antibody. As expected for a selective signal for autophagy, p62 showed a high degree of colocalization with ubiquitin. In contrast, ubiquitin signal was completely absent from Proteostat-positive protein aggregates (Figure 4H-I). We then examined whether the Proteostat-labeled structures corresponded to lysosomes, as has recently been reported (25–27). For this purpose, we co-immunostained cardiac sections of Eef1a2-cKO mice for LAMP1, a lysosomal membrane marker, together with p62 or Proteostat. Consistent with our previous observations, p62 did not colocalize with LAMP1, confirming that deletion of eEF1A2 impairs autolysosome formation (Figure 4J). In contrast, Proteostat-positive structures showed clear colocalization with LAMP1 (Figure 4K). These results indicate that protein aggregates in Eef1a2-cKO cardiomyocytes are associated with lysosomes and further demonstrate that the lack of p62–LAMP1 colocalization is not due to a deficiency in lysosome formation, but rather to a block in autophagic flux. Electron microscopy revealed largely unaltered sarcomere and mitochondrial morphology, and an accumulation of autophagosomes, but not autolysosomes in cardiomyocytes of Eef1a2-cKO versus control mice, again pointing towards a defect in autophagic flux due to eEF1A2 deletion (Figure 4L).

### eEF1A2 acts as chaperone in cardiomyocytes

Several studies have demonstrated a chaperone-like activity of eEF1A in vitro (28–30). To directly investigate whether increased levels of protein aggregates that we observed in our model might be due to a loss of this activity, we performed a luciferase-based protein folding assay. For this, we isolated cardiomyocytes from WT, Eef1a2-cKO, and Eef1a1/Eef1a2-cKO mouse hearts one month after Tamoxifen administration (Figure 5A). These cardiomyocytes were transduced with adenoviruses expressing firefly luciferase (under the SV40 promoter) or Renilla luciferase (under the thymidine kinase promoter) (31). Our results showed a significant reduction in luciferase activity in cardiomyocytes from Eef1a2-cKO and double cKO mice. More specifically, firefly luciferase activity decreased by ∼80% and Renilla luciferase activity by ∼40% compared to WT cardiomyocytes (Figure 5B, D). This reduction in luciferase activity was not due to differences in transcriptional activity, as luciferase mRNA levels were similar across all groups (Figure 5C, E). Additionally, protein synthesis appeared largely unaffected in Eef1a2-cKO cardiomyocytes (Figure 1G-H) and was decreased only slightly in the double cKO (Figure 2G, I). These results suggest that reduced translation alone does not account for the observed effects, rather implying reduced protein synthesis fidelity or defective protein folding as potential mechanisms. Since eEF1A has been implicated in maintaining protein synthesis fidelity (32, 33), we tested whether the decrease in luciferase activity might be due to increased amino acid misincorporation or stop codon read-through. To assess this, we transduced cardiomyocytes with adenoviruses encoding mutant firefly luciferase constructs carrying either the R216S substitution or a premature stop codon at position 210, along with WT Renilla luciferase as an internal control. As shown in Figure S11A-D, the activity of mutant firefly luciferase (R218S or STOP) was undetectable after normalization to Renilla, in knock-out similar to WT cardiomyocytes, indicating that amino acid misincorporation or read-through is unlikely to explain the reduction in luciferase activity. Accumulation of misfolded proteins has been shown to trigger the integrated stress response (ISR). We therefore analyzed phosphorylation of the eukaryotic initiation factor 2 α (eIF2α) in cardiac tissue, which is a common feature of the ISR. As shown in Figure 5F-H, we observed an increase in phospho-eIF2α as well as total eIF2α two months after eEF1A2 ablation.

**Figure 5.**
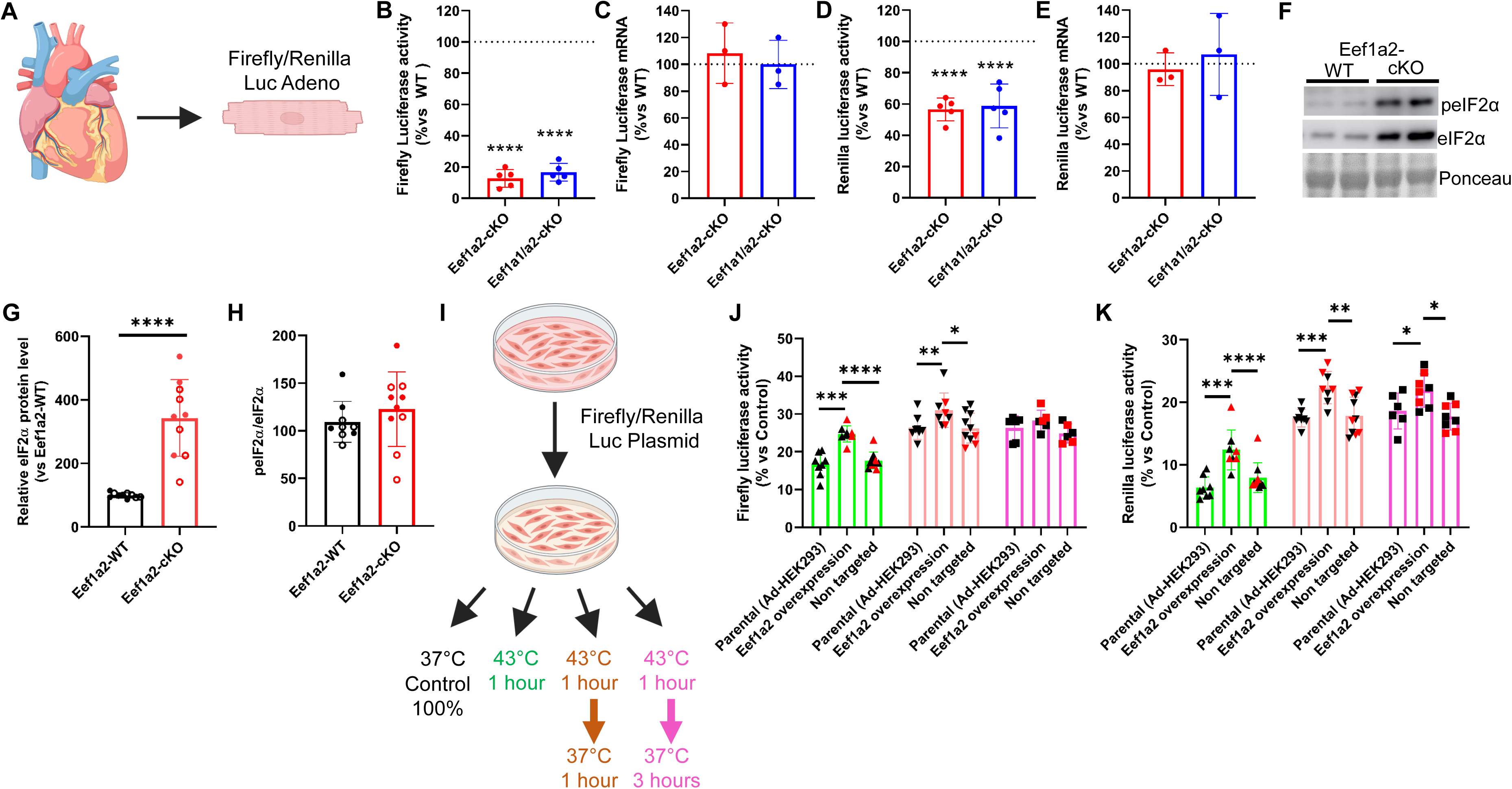
Effects of eEF1A2 on protein folding. (**A**) Schematic representation of the luciferase folding assay. Luc denotes luciferase. Adeno denotes Adenovirus. (**B**) Relative firefly luciferase activity measured in isolated cardiomyocytes from Eef1a2-cKO (n = 5) or Eef1a1/a2-cKO (n = 5) mice versus wild-type (WT). (**C**) Relative mRNA expression of firefly luciferase in isolated cardiomyocytes from Eef1a2-cKO (n = 3) or Eef1a1/a2-cKO (n = 3) mice versus WT. (**D**) Relative Renilla luciferase activity measured in isolated cardiomyocytes from Eef1a2-cKO (n = 5) or Eef1a1/a2-cKO (n = 5) mice versus WT. (**E**) Relative mRNA expression of Renilla luciferase in isolated cardiomyocytes from Eef1a2-cKO (n = 3) or Eef1a1/a2-cKO (n = 3) mice versus WT. (**F**) Representative Western blot analysis of the indicated proteins in cardiac tissue from Eef1a2-WT and Eef1a2-cKO mice 2 months after Eef1a2 deletion. (**G-H**) Quantification of total eIF2α expression (G) and the p-eIF2α/eIF2α ratio (H) in Eef1a2-WT (n = 10) and Eef1a2-cKO (n = 10) mice. ****P < 0.0001, unpaired two-tailed t-test. (**I**) Schematic representation of the protein unfolding/refolding assay. (**J**) Relative firefly luciferase activity measured in two independent Eef1a2-overexpressing clones (n = 4 each), two independent non-targeting control clones (n = 4 each), and parental cells (n = 8), normalized to their respective controls. *P < 0.05, **P < 0.01, ***P < 0.001, ****P < 0.0001; mixed-effects model followed by Tukey’s multiple-comparison test. (**K**) Relative Renilla luciferase activity measured in the same clones and parental cells as in (J), normalized to their respective controls. *P < 0.05, **P < 0.01, ***P < 0.001, ****P < 0.0001; mixed-effects model followed by Tukey’s multiple-comparison test. Filled circles denote males; open circles denote females.

Next, to investigate whether eEF1A2 functions as a molecular chaperone promoting protein folding, we generated a HEK293 cell line constitutively expressing eEF1A2 under the control of the CMV promoter using CRISPR/Cas9 technology. As shown in Figure S11E, we successfully isolated two independent clones that overexpress eEF1A2. Importantly, overexpression of eEF1A2 did not affect the expression levels of eEF1A1, which is the predominant paralog in HEK293 cells. We transfected the cells with a plasmid encoding both Firefly and Renilla luciferase and performed a luciferase unfolding/refolding assay (Figure 5I). Cells were subjected to a heat shock (43°C for one hour), and luciferase activities were measured immediately afterwards (unfolding assay), as well as after one and three hours of recovery (refolding assay). Our results showed that eEF1A2 overexpression partially protected both luciferases from heat-induced unfolding compared to WT or non-targeted control cells. Furthermore, the refolding of luciferase enzymes was accelerated in eEF1A2-overexpressing cells, with significant increased activity at one hour for both luciferases and at three hours for Renilla luciferase (Figure 5J-K).

Altogether, we demonstrate that eEF1A2 has a chaperone-like activity in vivo, whereby its loss leads to reduced folding efficiency of client proteins such as luciferase and further activates the ISR in cardiomyocytes.

### Rapamycin treatment partially rescues cardiomyopathy in Eef1a2-cKO mice

We hypothesized that defective autophagy with accumulation of autophagosomes and misfolded proteins might play a causative role in the development of cardiomyopathy in Eef1a2-cKO mice. Because increased mTORC1 activation as indicated by increased RPS6 phosphorylation was a potential reason for defective autophagic flux in these mice, we investigated whether pharmacological inhibition of mTORC1 might have beneficial effects. Treatment with the mTORC1 inhibitor Rapamycin was initiated one month after Tamoxifen administration (Figure 6A)—a time point at which cardiac function was still comparable between Eef1a2-cKO and WT mice, although eEF1A2 protein was already completely lost in the hearts of knock-out animals. As shown in Figure 6B, daily administration of Rapamycin for one month completely reversed the increased mortality in Eef1a2-cKO mice. Rapamycin also restored systolic cardiac function and reversed cardiac hypertrophy (Figure 6C-F). Interestingly, despite these improvements, cardiac fibrosis remained unaffected (Figure 6G). To better understand how Rapamycin improved cardiac function in Eef1a2-cKO mice, we analyzed myocardial protein synthesis, phosphorylated RPS6 abundance, autophagosomes, protein aggregates and fibrosis. As shown in Figure 6H-I, protein synthesis remained similar between the control and Rapamycin-treated groups. The abundance of representative 5’TOP mRNA encoded proteins (including RPS6, eEF2, and RPS15) as well as of ATG7 and Beclin1 were not significantly different between vehicle- and Rapamycin-treated Eef1a2-cKO hearts, despite complete loss of RPS6 phosphorylation following Rapamycin treatment (Figure 7A-C, S12A-D). In contrast, Rapamycin led to a significant reduction in p62 and LC3-I proteins in the heart. This effect was accompanied by a decrease in Proteostat and p62 positive cardiomyocytes compared to the vehicle-treated group (Figure 7D-E). Notably, we also observed a reduction in the expression of phospho-eIF2α and total eIF2α in response to Rapamycin, demonstrating that it relieves the ISR response in Eef1a2-cKO cardiomyocytes (Figure 7A-B).

**Figure 6.**
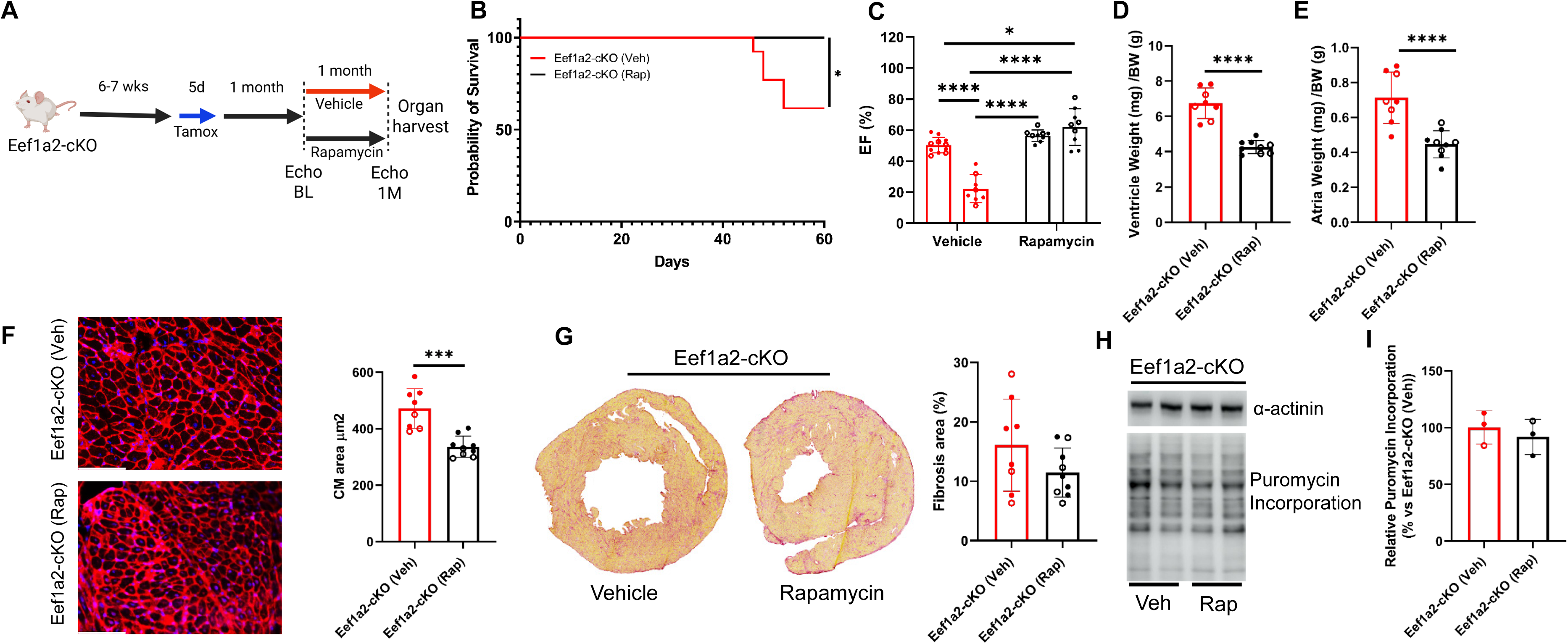
Effects of Rapamycin administration in Eef1a2-cKO mice. **(A)** Schematic representation of the experimental workflow. **(B)** Kaplan–Meier survival curves of vehicle-(n=13, 9/13 survived) or Rapamycin-treated (n=9, 9/9 survived) Eef1a2-cKO mice. *P < 0.05, log-rank (Mantel–Cox) test. **(C)** Left ventricular ejection fraction (EF) in Eef1a2-cKO mice treated with vehicle (red bars) or Rapamycin (black bars), measured before vehicle/Rapamycin administration (left bars, n = 11 and n = 9, respectively) and after 1 month of daily Rapamycin treatment (right bars, n = 8 and n = 9, respectively). ****P < 0.0001, two-way ANOVA followed by Tukey’s multiple-comparison test. (**D**) Ventricular weight relative to body weight in vehicle-(n = 8) or Rapamycin-treated (n = 9) Eef1a2-cKO mice. ****P < 0.0001, unpaired two-tailed t-test. **(E)** Atrial weight relative to body weight in vehicle-(n = 8) or Rapamycin-treated (n = 9) Eef1a2-cKO mice. ****P < 0.0001, unpaired two-tailed t-test. **(F)** Representative microscopic images of cardiac sections stained with wheat germ agglutinin (WGA) and corresponding quantification from vehicle-(n = 8) or Rapamycin-treated (n = 9) Eef1a2-cKO mice. ***P < 0.001, unpaired two-tailed t-test. **(G)** Representative Sirius Red staining of cardiac fibrosis and corresponding quantification from vehicle-(n = 8) or Rapamycin-treated (n = 9) Eef1a2-cKO mice. **(H)** Representative Western blot analysis of puromycin incorporation in cardiac tissue from vehicle- or Rapamycin-treated Eef1a2-cKO mice. **(I)** Quantification of puromycin incorporation in vehicle- (n = 3) or Rapamycin-treated (n = 3) Eef1a2-cKO mice. Filled circles denote males; open circles denote females.

**Figure 7.**
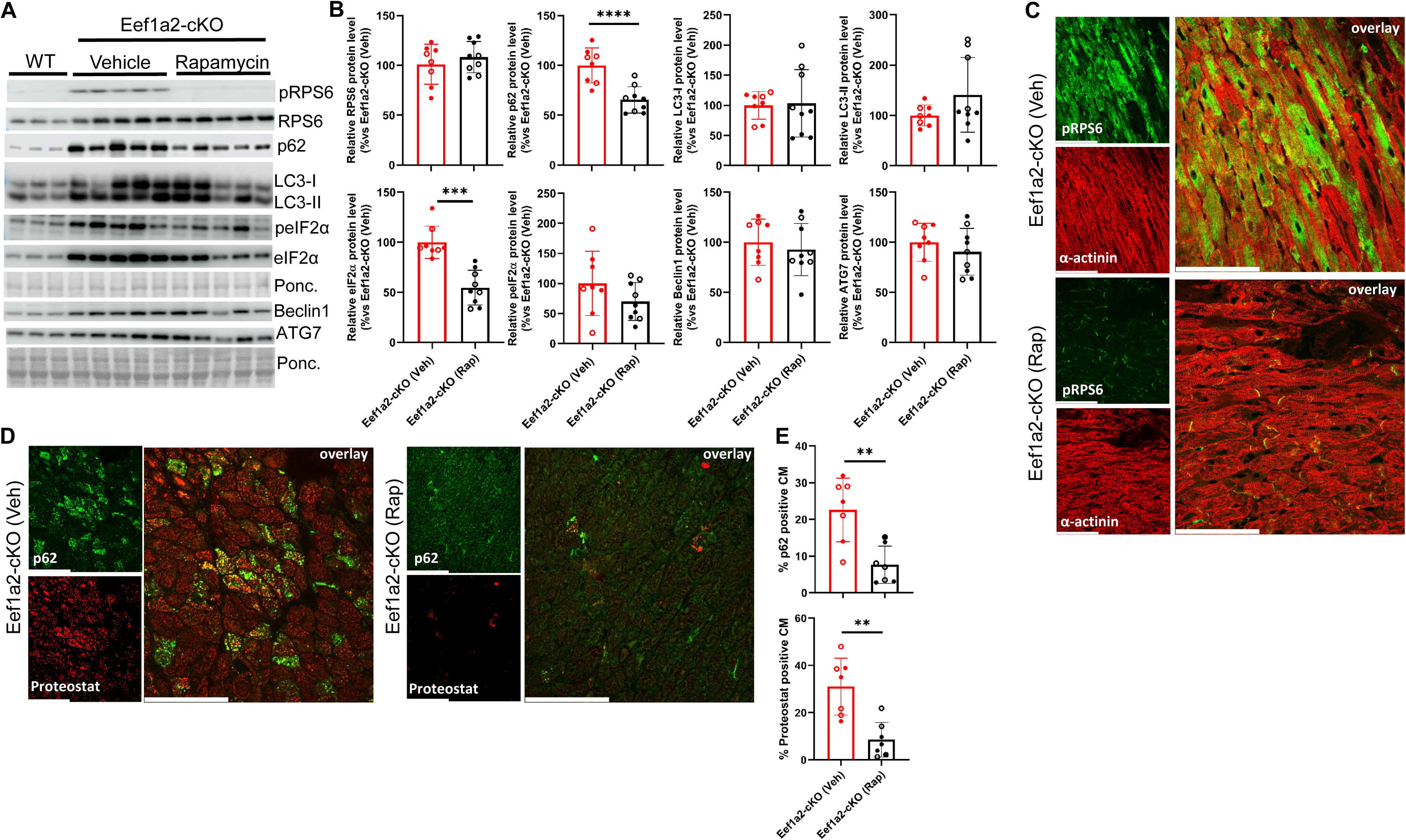
Effect of Rapamycin administration on altered proteostasis in Eef1a2-cKO mice. (**A**) Representative Western blot analysis of indicated proteins in cardiac tissue from vehicle- or Rapamycin-treated Eef1a2-cKO mice.(**B**) Quantification of protein expression shown in (A) in vehicle-treated (n = 8) or Rapamycin-treated (n = 9) Eef1a2-cKO mice. ***P < 0.001, ****P < 0.0001; unpaired two-tailed t-test.(**C-D**) Representative co-immunostaining of indicated proteins in cardiac tissue from vehicle- or Rapamycin-treated Eef1a2-cKO mice (scale bars, 100 µm). (**E**) Quantification of p62-positive (upper panel) Proteostat-positive (lower panel) and cardiomyocytes in cardiac tissue from vehicle-treated (n = 7) or Rapamycin-treated (n = 7) Eef1a2-cKO mice. **P < 0.01; unpaired two-tailed t-test.Filled circles denote males; open circles denote females.

In summary, our results indicate that the loss of eEF1A2 impairs cardiac proteostasis, contributing to cardiac dysfunction. Importantly, restoration of proteostasis by Rapamycin largely reversed the associated cardiac pathology in Eef1a2-cKO hearts.

## Discussion

One key observation from our study was the unaltered global protein synthesis activity in Eef1a2-cKO mice, despite the complete absence of eEF1A2 in cardiomyocytes. A previous report noted increased eEF1A1 protein expression in Eef1a2-cKO hearts; however, the underlying mechanisms and physiological significance were unknown (7). Here, we demonstrate for the first time that eEF1A2 regulates eEF1A1 at the post-transcriptional level, and that the upregulation of eEF1A1 serves as a compensatory mechanism. This is supported by our finding that double conditional knockout of both eEF1A1 and eEF1A2 led to mildly reduced global protein synthesis and 100% mortality approximately 6–7 weeks after gene deletion. Our results support the current paradigm that eEF1A1 and eEF1A2 expression is mutually exclusive, but also reveal a previously unrecognized layer of regulatory interaction between these two paralogs in the heart.

Protein synthesis varies significantly across organs and developmental stages. During postnatal cardiac development, it is markedly reduced, making adult cardiac tissue one of the organs with the lowest protein synthesis rates (34). Moreover, this downregulation coincides with the upregulation of eEF1A2 expression in the heart. This inverse correlation suggests that eEF1A2 may serve non-canonical functions in the cardiac context. Previous studies have shown that during translation, eEF1A remains associated with nascent polypeptides, and that it interacts with damaged proteins and facilitates their degradation via the proteasome (29, 35, 36). This suggested that eEF1A2 might play a role in protein quality surveillance by acting as a co-translational chaperone and in clearance of misfolded proteins. The protein aggregate formation and an mRNA-independent reduction in luciferase activity in cardiomyocytes lacking eEF1A2 argue in favor of this. Moreover, overexpression of eEF1A2 prevented protein unfolding and enhanced protein refolding during and after heat shock. Together, these findings demonstrate that eEF1A2 functions as a physiologically relevant co-translational chaperone in vivo. By doing so, eEF1A2 strongly promotes cellular proteostasis.

Two important questions remain unanswered: does eEF1A2 function as a direct chaperone, and does it exhibit specificity toward certain protein substrates? Regarding the first question, a recent study showed that eEF1A1, but not eEF1A2, transcriptionally regulates the expression of HSP70 and HSP27 after heat shock (37). Both molecular chaperones help cells cope with stress by preventing protein misfolding and aggregation. Conversely, several studies have demonstrated a direct role for eEF1A or eEF-Tu (Thermal unstable), the bacterial homologue of eEF1A, in the in vitro folding of several proteins, supporting a direct chaperone-like function (29, 30, 38, 39). Our results are more consistent with this latter possibility, as our Ribo-seq, RNA-seq, and proteomics analyses did not show activation of the heat-shock response. Nonetheless, we cannot fully exclude some contribution of HSPs in our model, especially since HSPs play a pivotal role in heart failure (40). For the second question, a previous study (29) found that eEF1A associates with nascent polypeptides independently of the Nascent Polypeptide-Associated Complex (NAC). This suggests that client proteins of eEF1A may differ from those bound by NAC, supporting the idea that eEF1A could possess its own set of specific substrates. Our findings reveal for the first time that eEF1A2 also regulates the expression of ribosomal subunits and elongation factors at the post-transcriptional level. The mRNAs encoding these proteins are characterized by a 5′ TOP motif (17). mTORC1 is a protein complex that regulates several essential cellular processes, including autophagy and protein synthesis, functioning as a key nutrient and energy sensor (41). It is also well-established that mTORC1 activation enhances the translation of 5′ TOP mRNAs (19–21, 42). The observed increase in translation efficiency of 5′ TOP mRNAs and the elevated levels of phosphorylated RPS6, a surrogate of mTORC1 activity, support a role for mTORC1 in this regulatory mechanism. However, treatment with Rapamycin, a well-known mTORC1 inhibitor, neither affected the expression of 5′ TOP mRNAs nor global protein synthesis in our model, although it strongly improved cardiac function and survival. Together, our results suggests that the elevated 5′ TOP mRNA levels may help to maintain protein synthesis in the absence of eEF1A2, but do not account for the development of cardiomyopathy. Notably, 5′ TOP mRNA translation in adult cardiomyocytes is mTORC1-dependent, but Rapamycin-resistant in vivo, as has also been reported in other cell types cultured in vitro (43).

Activation of the ISR inhibits global translation (44) through phosphorylation of eIF2α (45), in order to restore protein homeostasis (46). Our results clearly demonstrate ISR activation in Eef1a2-cKO cardiac tissue, indicated by increased eIF2α phosphorylation (47). However, this activation occurs without a corresponding decrease in global protein synthesis. This intriguing finding may be explained by the observed increase in ribosomal proteins and translation factors. Such compensatory upregulation could help to maintain protein synthesis even in the presence of ISR activation. The significance of the increased levels of both phosphorylated and total eIF2α in Eef1a2-cKO cardiomyocytes remains unknown. Of note, a recent study demonstrated that phosphorylation of eIF2α by HRI exerts a protective role in a model of mitochondrial cardiomyopathy (48). Whether this response is ultimately protective or detrimental remains an open question that future studies will need to address.

Autophagy is a highly conserved pathway in eukaryotes and is essential for maintaining cellular homeostasis by degrading excess or damaged intracellular components (49). There are three main types of autophagy: macroautophagy, microautophagy, and chaperone-mediated autophagy (CMA) (50). The relationship between macroautophagy, involved in the clearance of protein aggregates and entire organelles, and cardiac dysfunction following cardiac stress, is complex and – at times – contradictory. For example, Kuhn et al. (51) and Maejima et al. (52) demonstrated that overexpression of FYCO1 or Mst1, both regulators of the autophagy machinery, are essential for adaptation to cardiac stress. Similarly, Nakai et al. (53) showed that inhibition of autophagy through Atg5 deletion worsens cardiac function under stress conditions. In contrast, Zhu et al. (54) reported that constitutive overexpression of Beclin1, a key regulator of early autophagosome formation, exacerbates cardiac dysfunction following transverse aortic constriction (TAC), highlighting the delicate balance required in autophagy regulation.

Our findings indicate that macroautophagy is markedly impaired in eEF1A2-deficient hearts and that this defect is closely associated with the onset of cardiac dysfunction. The accumulation of p62 and LC3-II, together with the presence of p62/ubiquitin-positive structures, suggests a blockade in autophagic flux rather than enhanced autophagosome formation (24). Consistent with this interpretation, electron microscopy analysis reveals a predominance of autophagosomes over autolysosomes, indicating defective autophagosome maturation or clearance. Moreover, the lack of colocalization between p62-positive bodies and the lysosomal marker LAMP1 further supports the conclusion that cargo delivery to, or degradation within lysosomes is compromised in eEF1A2-deficient cardiomyocytes. Importantly, we demonstrate that mTORC1 activation, which inhibits macroautophagy (55), is a key driver of this phenotype, as Rapamycin, a specific mTORC1 inhibitor, restores macroautophagy activity and improves cardiac function in eEF1A2-deficient hearts. Our results support the concept that autophagy plays a critical role in maintaining cardiac function and emphasize the importance of autophagy dysfunction as shared pathogenic mechanism across different genetic models of cardiac dysfunction. Despite the clear relationship between eEF1A2, mTORC1 activation and impaired macroautophagy, the precise molecular mechanism remains unknown. Based on our findings, we propose two possible scenarios. In the first, the absence of eEF1A2 directly activates cardiac mTORC1, leading to a subsequent reduction in autophagy activity and subsequent accumulation of autophagosomes and misfolded proteins (56). How the lack of eEF1A2 leads to increased mTORC1 activation remains unclear, but may involve sensing of increased nutrient availability due to increased levels of aminoacyl-tRNAs, or a stimulating effect on the assembly of the mTORC1 complex. A previous study from Qian et al. (57) demonstrated that intracellular chaperones act as inhibitors of mTOR complex assembly and showed that partial downregulation of HSP90 was sufficient to activate mTORC1. Here, we demonstrate that eEF1A2 acts as a chaperone-like protein in cardiomyocytes and may function similarly to HSP90 in regulating mTORC1 activity. Notably, we observed increased expression of ATG7 and Beclin1, two key activators of autophagosome formation, along with an accumulation of autophagosomes but a lack of autolysosomes in hearts of Eef1a2-cKO mice. Together, these findings suggest a defect in autophagosome–lysosome fusion in these mice. In the second scenario, the accumulation of unfolded proteins may sequester lysosomes away from autophagosomes. This hypothesis is supported by the extensive presence of p62-positive bodies and protein aggregates within the same cardiomyocytes and, more importantly, by the colocalization of LAMP1 with protein aggregates but not with p62-positive bodies. In this context, inhibition of mTORC1 by Rapamycin may reduce the accumulation of misfolded proteins either directly (58) or indirectly, by improving the fidelity of protein translation (59) or by increasing ubiquitination and proteasomal proteolysis (60).The two scenarios are not mutually exclusive and may synergize in driving mTORC1 activation in the absence of eEF1A2.

Among our main findings are the positive effects of Rapamycin administration on cardiomyopathy in Eef1a2-cKO mice. Rapamycin and derivatives are currently approved immunosuppressants to prevent organ rejection after transplantation and to reduce re-stenosis in vessels for example after insertion of a coronary artery stent. Our study expands the spectrum of cardiac disease models in which Rapamycin may exert therapeutic benefits. Previous studies, such as those by Choi et al. and Ramos et al. (61, 62), have shown that Rapamycin and its derivatives prevent cardiomyopathy in laminopathies. Moreover, Rapamycin administration has been reported to prevent or reverse cardiac dysfunction following transverse aortic constriction (TAC) in mice (63, 64). In addition, Rapamycin increases lifespan and delays age-related dysfunction in different organism including rodents (65–69) and flies (59). Here, we demonstrate that administration of Rapamycin in the early stage of Eef1a2-cKO cardiomyopathy reverses systolic dysfunction and cardiac hypertrophy, and, most importantly, improves mouse survival. Notably, however, Rapamycin did not reverse fibrosis.

This observation is consistent with previous findings in Lamin A/C–mediated cardiomyopathies, where Rapamycin restored cardiac function, but did not reduce fibrosis (61). While it might seem counterintuitive to treat cardiomyopathy caused by lack of a canonical translation elongation factor with an mTORC1 inhibitor, which may result in reduced protein synthesis, our study emphasizes the therapeutic effectiveness of Rapamycin as an enhancer of autophagic flux, while not effecting the overall protein synthesis rate in our disease model. While our study clearly shows short-term benefits of Rapamycin treatment on cardiac function and survival, its long-term effects beyond two months after eEF1A2 ablation remain uncertain. Future experiment will address this important question, in order to assess whether mTORC1 inhibition represents a tailored treatment option for patients suffering from cardiomyopathy due to mutations in the *EEF1A2* gene.

In summary, we characterize eEF1a2 as a critical mediator of cardiac proteostasis, independent of its canonical role in protein synthesis. Mechanistically, we demonstrate that eEF1A2 deficiency impairs cardiac function by reducing both autophagic activity and protein folding capacity, and uncover mTORC1 inhibition as an effective strategy to restore cardiac proteostasis and function in Eef1a2-cKO hearts. While our study focuses on the cardiac-specific role of eEF1A2, this paralogue is also highly expressed in neurons, explaining why mutations in *EEF1A2* are not only associated with cardiomyopathy, but also with intellectual disability, intractable epilepsy, and, in some cases, autism. It will be of great interest to explore whether some of these patient-derived eEF1A2 mutations also impair proteostasis in the brain, and if mTORC1 inhibitors have therapeutic potential beyond the cardiological context for these severe neurological symptoms.

## Methods

### Sex as a biological variable

Male and female mice were used in all experiments, except for Ribo-Seq and polysome profiling, where only male mice were used to reduce variability.

### Mouse lines

The *Eef1a1* and *Eef1a2* conditional knockout mouse lines were generated by Cyagen Biosciences (Santa Clara, CA, USA). *Eef1a1* and *Eef1a2* conditional knock-out mouse lines were generated by flanking exons 2–8 (Eef1a1 flox/flox) or exons 2–5 (Eef1a2 flox/flox) with loxP sites. Floxed mice were crossed with heterozygous αMHC–MerCreMer transgenic mice to obtain cardiomyocyte-specific, Tamoxifen-inducible gene deletion. For double *Eef1a1* and *Eef1a2* conditional knockout mice, we first generated Eef1a1 flox/flox; Eef1a2 flox/flox mice, which were the crossed with αMHC–MerCreMer +/-transgenic mice to obtain cardiomyocyte-specific, Tamoxifen-inducible gene deletion.

At least three independent cohorts (derived from different breeding pairs) were used for all in vivo experiments and for experiments using isolated cardiomyocytes.

### Tamoxifen injection

A stock solution of Tamoxifen (8 mg/mL) was prepared in Miglyol® 812 (Caesar & Loretz GmbH, Hilden, Germany) and heated to 56 °C until completely dissolved. Mice aged 6–7 weeks received Tamoxifen at a dose of 40 µg/g body weight via daily intraperitoneal injections for five consecutive days.

### Rapamycin administration

A stock solution of Rapamycin was prepared at 1.5 mg/mL in a vehicle containing 5% Tween 80 (Sigma) and 5% PEG-400 (Sigma) as previously described (62). Vehicle control solutions were prepared identically but without Rapamycin. Rapamycin was administered daily by intraperitoneal injection at a dose of 8 mg/kg body weight.

### Transthoracic echocardiography

Echocardiography was performed under 1% isoflurane anesthesia as previously described using a 30– 40 MHz linear transducer (MX-550D) with a Vevo 3100 imaging system (VisualSonics, Toronto, Canada) (70). Mice were placed in the supine position on a heated platform to maintain body temperature. Data were analyzed using Vevo Lab software (v5.5.0). All echocardiographic analyses were performed in a blinded manner, with the operator unaware of mouse genotype and treatment. Parameters were recorded in B-mode and M-mode in both parasternal long-axis (PSLAX) and short-axis view at the papillary muscle level. The change of LV diameter length from end-diastole to end-systole was used to calculate LV ejection fraction and fractional shortening, which were measured with the PSLAX mode (70).

### Organ harvest

At the end of the experiment, mice were weighed and euthanized by cervical dislocation to collect different organs. Organs were washed in cold PBS to remove the blood, and additionally the heart was washed in 0.5% (w/v) KCl in PBS. The heart was transversely cut into half, and the upper half was immediately embedded in optimal cutting temperature (OCT) compound for histological analysis, while the left ventricle from the remaining half was dissected and snap-frozen in liquid nitrogen.

### RNA isolation and quantitative PCR

For mRNA isolation, cardiac tissue or cultured cells were homogenized in 1 mL TRIzol reagent. Cardiac tissue was homogenized using a TissueLyser II (Qiagen) for 2 minutes at 20 Hz. Following centrifugation at 12,500 × g for 10 minutes at 4 °C, 200 µL chloroform was added, and samples were incubated for 10 minutes. Samples were then centrifuged at 12,000 × g for 15 minutes at 4 °C, and the aqueous phase was collected. RNA was precipitated by adding 500 µL isopropanol and incubation for 30 minutes on ice, followed by purification using NucleoSpin RNA columns (Macherey-Nagel) according to the manufacturer’s instructions.

cDNA was synthesized using the Maxima H Minus First Strand cDNA Synthesis Kit (Thermo Fisher Scientific, K1652). Quantitative PCR (qPCR) was performed using PowerUp™ SYBR™ Green Master Mix with ROX as a reference dye (Thermo Fisher Scientific, A25742) on an AriaMx Real-Time PCR System (Agilent, G8830A). All qRT-PCR primer sequences are listed in Supplemental Table S7.

### Protein isolation and Western blotting

For protein extraction, cardiac tissue or cells were lysed in sampling buffer (30 µL/mg tissue) containing 50 mM Tris-HCl (pH 6.7), 2% SDS, 10% glycerol, a protease inhibitor cocktail (Roche), and phosphatase inhibitor cocktail set V (Millipore). Western blotting was performed using standard procedures. All used antibodies and their dilutions are listed in Supplemental Table S8.

### Histology

OCT-embedded organs were sectioned using a cryostat at a thickness of 7 µm for immunofluorescence and 12 µm for Picrosirius red staining. For immunofluorescence, tissue sections were fixed in 4% paraformaldehyde (PFA), permeabilized with 0.3% Triton X-100 in PBS, and blocked with 3% BSA. Primary and secondary antibodies used in this study are listed in Supplemental Table S8. Samples were visualized using a Leica SP8 confocal microscope (Leica Microsystems, Wetzlar, Germany), and images were analyzed using Leica Application Suite X (LAS X) software version 3.7. Picrosirius red staining was performed by fixing tissue sections in 100% ice-cold acetone, followed by staining according to a standard protocol. Tissue sections were scanned using a Zeiss Axio Scan.Z1 system (Carl Zeiss Microscopy GmbH, Jena, Germany), and images were analyzed using ZEN 2.6 Blue Edition software (Carl Zeiss Microscopy GmbH, Jena, Germany).

### Proteostat staining of cardiac tissue

Tissue sections were fixed in 3.7% methanol-free formaldehyde (Pierce) diluted in 1× assay buffer for 30 minutes, washed twice with phosphate-buffered saline (PBS), and permeabilized with 0.5% Triton X-100 in 1× assay buffer for 30 minutes. After two additional washes, sections were incubated with PROTEOSTAT® Aggresome Detection Reagent (Enzo Life Sciences) diluted in assay buffer according to the manufacturer’s instructions for 30 minutes. Next, sections were co-immunostained for selected proteins as described above.

### Adenovirus generation and infection

Adenoviral vectors were used to overexpress standard, STOP, and R218S firefly luciferase together with Renilla luciferase. To eliminate a PacI restriction site while preserving the original amino acid sequence, a silent mutation was introduced at leucine 440 of firefly luciferase (TTA→TTG) in the original plasmid (kindly provided by Dr. Xie, (31)). The modified luciferase sequences were subcloned into the promoterless pShuttle vector (Addgene plasmid #16402), and recombinant adenoviruses were generated using the AdEasy Adenoviral Vector System (Agilent Technologies, Waldbronn, Germany) in AD-293 cells (Agilent) according to the manufacturer’s instructions. Viral titers were determined using the AdEasy Viral Titer Kit (Agilent Technologies). For adenoviral infection, cardiomyocytes were maintained in medium containing heat-inactivated fetal calf serum.

### Luciferase assay

100 000 isolated cardiomyocytes were plated per well (of a 6 well-plate) treated with laminin A (Santa Cruz). After 3 hours, cells were infected overnight at a multiplicity of infection (MOI) of 100. Luciferase activity (Firefly and Renilla) was measured after 48 hours post infection using the Dual-Luciferase® Reporter Assay System (Promega) according to the manufacturer’s instructions. For each isolation, signals from non-infected cardiomyocytes were used as background control.

### Electron microscopy

Immediately after cervical dislocation, hearts were perfused with freshly prepared Perfixol solution containing 24 mM cacodylic acid (Sigma, Cat. No. 20835), 4% paraformaldehyde, and 2.5% glutaraldehyde (Sigma, Cat. No. 354400), pH 7.2. The left ventricle was then dissected and cut into small squares, which were incubated overnight at 4 °C in Perfixol. After rinsing in buffer, the samples were further fixed in 1% osmium in 0.1M cacodylate buffer for 1h, washed in water, and incubated in 1% uranyl acetate in water overnight at 4°C. Dehydration was done in 10 min steps in an acetone gradient followed by stepwise Spurr resin infiltration at room temperature and polymerization at 60°C. The blocks were sectioned using a Leica UC6 ultramicrotome (Leica Microsystems Vienna) in 70 nm thin sections and placed on formvar-coated slot grids, post-stained with 3% uranyl acetate in water and Reynold’s lead citrate, and imaged on a JEOL JEM-1400 electron microscope (JEOL, Tokyo) operating at 80 kV and equipped with a 4K TemCam F416 (Tietz Video and Image Processing Systems GmBH, Gautig).

### Ribo-seq analysis

Left ventricular tissue was homogenized in 500 µL ice-cold polysome buffer (20 mM Tris-HCl, pH 7.4, 10 mM MgCl₂, 200 mM KCl, 2 mM DTT, 1% Triton X-100, 1 U DNase/µL) supplemented with 100 µg/mL cycloheximide using a tissue homogenizer. Samples were centrifuged at 20,000 × g for 15 min at 4°C, and the supernatant was immediately processed. An aliquot (100 µL) was reserved for total RNA extraction using TRIzol, while ribosomal footprints were isolated from the remaining lysate using MicroSpin™ S-400 HR columns (Merck) according to the manufacturer’s instructions. SDS (10%, 10 µL per 100 µL eluate) was added, and RNA was purified using the RNA Clean & Concentrator-25 kit (Zymo Research).

Total RNA (5 µg) was subjected to rRNA depletion using the RiboPool Kit (Biozym) and precipitated with GlycoBlue, NaCl and isopropanol. Fragmentation was performed by incubating RNA in 2× fragmentation buffer (2 mM EDTA, 85 mM NaHCO₃, 120 mM Na₂CO₃) at 95°C for 12 min. Fragmented RNA was size-selected (25–35 nt) by 15% TBE–urea PAGE, visualized with SYBR Gold (Invitrogen), and gel slices were eluted overnight at 4°C in 0.3 M NaCl containing RNaseOUT. RNA was recovered by filtration and isopropanol precipitation. End repair was performed using T4 polynucleotide kinase (NEB) at 37°C for 1 h, followed by RNA precipitation and resuspension in RNase-free water.

Small RNA libraries were prepared using the NEXTflex Small RNA-Seq Kit v3 (PerkinElmer) with pre–size-selected RNA. Adapter-ligated libraries were amplified using a qPCR-determined number of cycles to avoid overamplification. Library quality and size distribution were assessed using a Bioanalyzer High Sensitivity DNA kit (Agilent). Sequencing was performed on an Illumina NextSeq 550 using the NextSeq 500/550 High Output Kit v2.5 (75 cycles).

Raw reads from four WT, four Eef1a2-cKO, and four Eef1a1/Eef1a2-cKO biological replicates were processed by trimming adapters with Cutadapt and removing rRNA/tRNA-mapping reads using Bowtie2. Remaining reads were aligned to the mouse reference genome (Ensembl release 98) using STAR v2.7.3a, retaining only uniquely mapped reads. Quality control and P-site assignment were performed using Ribo-seQC. All samples exhibited robust three-nucleotide periodicity (81–84%), with predominant read lengths of 28–29 nt. P-sites overlapping annotated coding sequences were quantified to calculate codon frequencies, and relative usage differences between genotypes were determined.

### RNA-seq and bioinformatics

Quality control of RNA samples (Agilent 2100 Fragment Analyzer) and library preparation (DNBSEQ Eukaryotic Strand-specific mRNA library) were performed by BGI, Hong Kong. Bulk RNA-seq from all samples was performed with single-end 50 base pair sequence read length by BGI with an Illumina HiSeq 2500. Initially, the trimming of adapter sequences from fastq files was performed with the R package FastqCleaner. For aligning the reads to the reference genome (mm10) after trimming, the R package bowtie2 alignment tool was used. Gene annotation was performed with the bioMaRt R package. Library size of the samples was normalized to counts per million (cpm) and transformed into log2 values followed by calculation of differential gene expression with the edgeR package of R. Significant changes in gene expression between compared groups were filtered based on FDR < 0.05 and fold change > 1.0. Gene ontology analysis was performed with Metascape and DAVID online tools.

### Ribosome profiling

Tissue isolation for polysome profiling: Quickly after euthanizing the mice, hearts were perfused through the apex with a solution containing PBS and cycloheximide (150 µg/ml), for stalling the ribosomes. Next, the heart was removed and washed in PBS/CHX solution to remove the remaining blood, and snap frozen in liquid nitrogen.

### Tissue lysis for polysome profiling

The left isolated left ventricle was lysed in 900 µL ice-cold polysome buffer without Triton X-100 (20 mM Tris-HCl, pH 7.4, 10 mM MgCl₂, 200 mM KCl, 2 mM DTT, 1 U DNase/µL) supplemented with 100 µg/mL cycloheximide using the TissueLyser II (Qiagen) for 2 minutes at 200 Hz. Next, Triton X-100 was added to reach 1% and samples were incubated for 30 minutes at 4 °C in an endless end to end rotor followed by centrifugation at 17,000 g for 6 minutes at 4 °C. The supernatant was collected and centrifuged again as before.

### Polysome fractionation and profiling

250 µl of lysate per gradient was loaded on a sucrose gradients (17.5 % - 50 %) and centrifuged for 2 hours (40 000 rpm at 4°C) in a SW60 rotor (Acceleration: 7, Deceleration: 1). Next, gradients were eluted with a Teledyne Isco Foxy Jr. system into 16 fractions of similar volume and the optical density of the fractions was measured.

### mRNA and protein isolation

Half of the volume of each fraction was used for mRNA isolation whereas the other half was subjected to protein isolation. For mRNA isolation, 1 mL of Trizol was added to each fraction and the mRNA was isolated as previously described. For protein isolation we used a method previously described by Poetz et al (71).

### Isolation and cultured of adult cardiomyocytes

Isolation and culture of adult mice cardiomyocytes were done as previously described by Acker-Johnson et al. (72) with minor changes as suggested by Callagham et al (73).

### Generation of stable overexpressing Eef1a2 HEK lines

Mouse Eef1a2 was cloned and inserted in pAAVS1-Nst-MCS. Next, AD-293 cells were transfected with pAAVS1-Nst-Eef1a2 (74) and pX459-HypaCas9-AAVS1_sgRNA (75). 48 hours later, transfected cells were selected by puromycin incubation for 72 hours and single cells were grown. Two different single clones that overexpressed Eef1a2 were selected. Two single clones without Eef1a2 overexpression and parental cells were used as control.

### Luciferase unfolding and refolding assay

AD-293 cells parental cells, as well as clones overexpressing Eef1a2 and non-targeted Eef1a2 control clones were transfected with the luciferase plasmids described in the Adenovirus section using Lipofectamine 3000 (Invitrogen). The following day, cells were split and seeded at a density of 9 × 10⁵ cells per well. On the next day, cycloheximide (CHX; final concentration 20 µg/mL) and MOPS buffer (pH 7.0; final concentration 20 mM) were added to the culture medium and incubated for 30 minutes. Cells were then subjected to heat shock at 43 °C for one hour or maintained at 37 °C as control. After heat shock, cells were allowed to recover at 37 °C for either one hour or three hours. At the indicated time points, cells were lysed with 400 µL of 1× Passive Lysis Buffer (Promega), and Firefly and Renilla luciferase activities were measured using the Dual-Luciferase® Reporter Assay System (Promega).

### Proteomic analysis

Sample preparation, mass spectrometry, and proteomic analyses were performed as previously described (76). Protein changes between conditions were classified as significant if they met an FDR < 20% with a fold change ≥ 50%. Only significantly changed proteins were included in the analysis. Pathway enrichment analysis was performed using the Metascape online platform.

### Metabolomics analysis

Frozen sample material was extracted with 360 µL 100% methanol containing 10 µL 0.02 mg/ml Ribitol for 15 min at 70°C with vigorous shaking. After the addition of 200 µL 100% chloroform, samples were shaken at 37°C for 5 min. To separate polar and organic phases, 400 µL HPLC-grade water were added and samples were centrifuged for 10 min at 11,000x g and 350 µL of the polar (upper) phase were transferred to a fresh glass vial and dried using a vacuum concentrator (Eppendorf Concentrator Plus) without heating.

### Derivatization (Methoximation and Silylation)

Sequential on-line methoximation and silylation reactions were performed using an AOC-6000plus autosampler (Shimadzu). Methoximation was performed by adding 20 µL 20 mg/ml methoxyamine hydrochloride (Sigma 226904) in pyridine (Sigma 270970) and incubation at 37°C for 90 min at 250 rpm. For silylation reactions, 45 µl of N-methyl-N-(trimethylsilyl) trifluoroacetamide (MSTFA; Sigma 69479) were added and samples were incubated at 37°C for 30 min with gentle shaking. Before injection, samples were incubated at room temperature for 2 hours.

### Gas Chromatography/Mass Spectrometry (GC/MS) analysis

For GC/MS analysis, a Shimadzu GC-MS/MS-system (TQ8040-NX) was used fitted with a Rxi-5Sil MS column (30 meter x 0.25 mm x 0.25 µm; Restek Corporation, Bellefonte). The GC was operated with an injection temperature of 280°C and 1 µL sample was injected with a split of 60. The GC temperature program started with a 1 min incubation at 65°C followed by a 6°C/min ramp up to 220°C, a 20°C/min ramp up to 330°C and a bake-out at 330°C for 5 min using helium as carrier gas with constant linear velocity. The mass spectrometer was operated with the ion source at 200°C, the interface at 280°C and a solvent cut time of 4.5 min. Data recording was in MRM mode using the cell specific selection of the Shimadzu Smart Metabolite Database. The Shimadzu LabSolution Insight software was used for data processing. Only metabolites detected in at least 4 of 5 biological replicates were included in the analysis. Metabolites showing significant differences relative to the control group were identified using one-way ANOVA. Pathway enrichment analysis was performed using the MetaboAnalyst 6.0 online platform.

### Schematic representations

Schematic representations were prepared with Biorender software.

### Statistics

All statistical analysis was performed using GraphPad Prism 8. Graphs are presented as mean ± SD. Each statistical test is included in the respective Figure legend. P values less than 0.05 were considered statistically significant.

### Study approval

The Regional Council Karlsruhe approved all animal studies under the protocol 35-9185.81/G-92/20.

## Data availability

All data associated with this study are present in the paper or Supplementary Materials.

## Author contributions

M.-G. conceptualized the study, designed and performed most of the experiments, analyzed data and wrote the first draft of the manuscript. N.W., J.E., S.R., F.S., V.S., A.G., M.K., N.W., E.H., F.A.T., S.H., M.B., M.R. performed experiments. J. R.-O., J.C., G.P., F.S. and E. A. R.-Z. analyzed data. N.F., G.S., M.V., N. H. and G.D. supervised experiments and critically revised the manuscript. J.H. initiated, designed and conceptualized the study, supervised experiments and prepared the final draft of the manuscript. All authors read and approved the manuscript.

## Acknowledgements

The authors are grateful to the Core Facilities Pre-Clinical Models and Live Cell Imaging Mannheim (LIMA), as well as for the support provided by the Electron Microscopy Core Facility (EMCF; RI_00565) at Heidelberg University. The authors gratefully acknowledge the data storage service SDS@hd supported by the Ministry of Science, Research and the Arts Baden-Württemberg (MWK) and the German Research Foundation (DFG) through grant INST 35/1803-1 FUGG and INST 35/1804-1 LAGG.

---

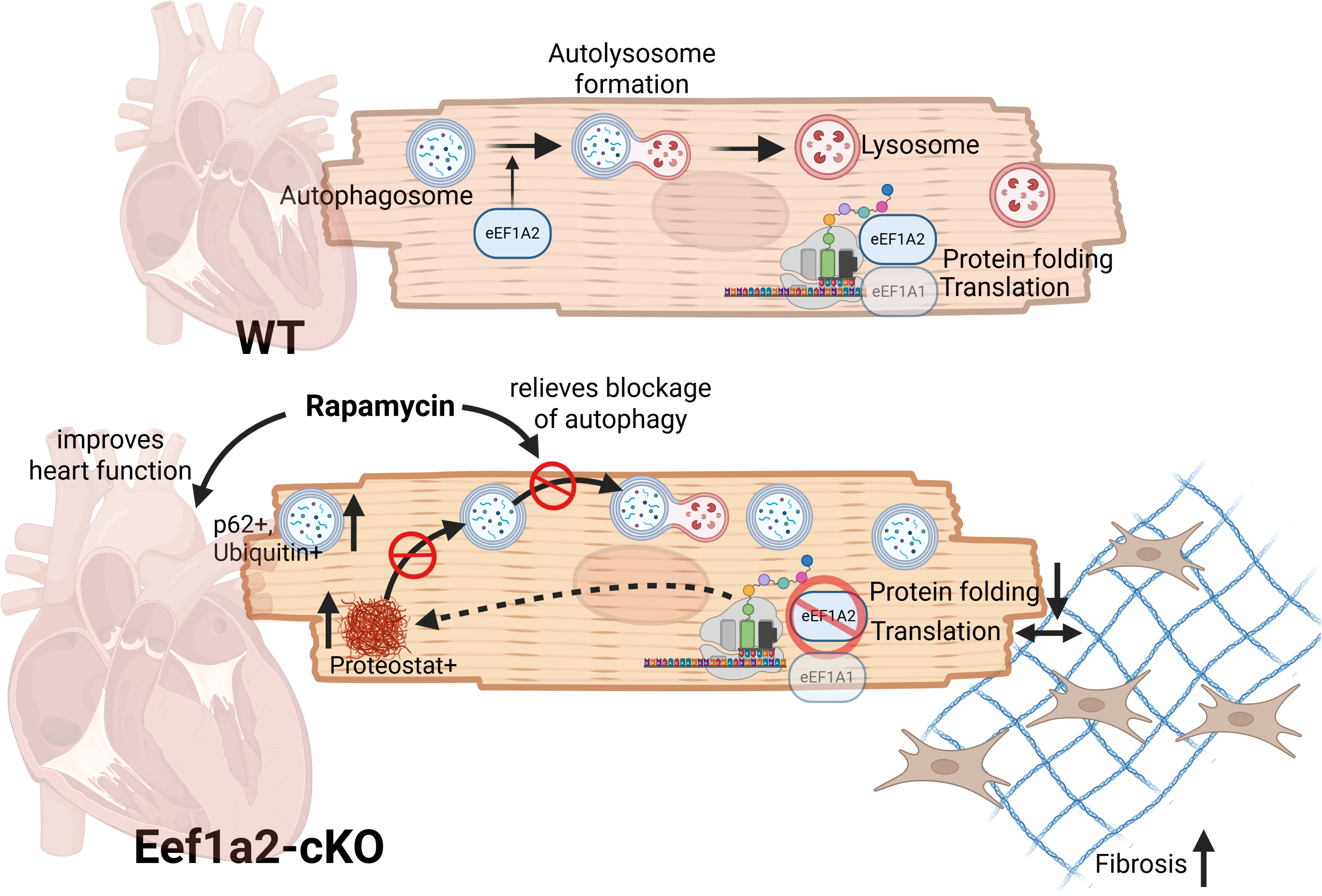

## SUPPLEMENTAL MATERIAL

**Table S1:** Echocardiography and morphometry two months after Tamoxifen administration.

|  | Male |  | Female |  |
| --- | --- | --- | --- | --- |
|  | Eef1a2-WT | Eef1a2-cKO | Eef1a2-WT | Eef1a2-cKO |
| **Echo** | n=5 | n=5 | n=5 | n=5 |
| BPM | 453.7±11.81 | 490.6±14.75 | 462±12.27 | 448±13.81 |
| EF (%) | 43.26±3.912 | 20.28±1.052*** | 41.88±5.329 | 16.53±5.164** |
| IVS; d (mm) | 1.13±0.032 | 1.025±0.048 | 1.09±0.062 | 0.976±0.027 |
| IVS; s (mm) | 0.823±0.022 | 0.864±0.035 | 0.7473±0.047 | 0.818±0.008* |
| LVID; d (mm) | 4.133±0.064 | 4.504±0.15* | 4.054±0.085 | 3.965±0.111 |
| LVID; s (mm) | 3.332±0.077 | 4.041±0.21** | 3.139±0.092 | 3.570±0.074** |
| LVPW; d (mm) | 0.757±0.024 | 0.676±0.023* | 0.7379±0.048 | 0.693±0.029 |
| LVPW; s (mm) | 0.983±0.031 | 0.752±0.035**** | 1.021±0.058 | 0.807±0.038** |
| **Morphometry** | n=5 | n=5 | n=5 | n=5 |
| BW (g) | 26.91±0.80 | 26.89±0.77 | 22.04±0.86 | 21.23±0.53 |
| VW/BW (mg/g) | 4.613±0.3318 | 5.446±0.148* | 4.806±0.117 | 5.514±0.349 |
| AW/BW | 0.1921±0.013 | 0.543±0.021**** | 0.213±0.012 | 0.525±0.072** |
| Lu/BW | 4.963±0.13 | 6.233±0.25** | 6.146±0.099 | 6.761±0.450 |

Data are shown as mean ± SD. \*p<0.05, \*\*p<0.01; \*\*\*p<0.001; \*\*\*\*p<0.0001 vs. WT at the same time point. Abbreviations: Echo= Echocardiography; BPM= beats per minute; EF= ejection fraction; IVS= interventricular septum; d= diastolic; s= systolic; LVID= left ventricular internal diameter; LVPW=left ventricular posterior wall thickness; BW= body weight; VW= ventricular weight; AW=atrial weight; Lu= lung weight.

**Table S2:** Echocardiography and morphometry four months after Tamoxifen administration.

| | Wild-type (WT) | αMHC-MerCreMer |
| --- | --- | --- |
| **Echo** | n=9 | n=8 |
| BPM | 459.8±15.05 | 448.8±12.27 |
| EF (%) | 49.94±3.649 | 54.36±3.656 |
| IVS;d (mm) | 0.8580±0.04371 | 0.7927±0.04605 |
| IVS;s (mm) | 1.211±0.06752 | 1.144±0.07796 |
| LVID;d (mm) | 3.887±0.07556 | 3.622±0.07481* |
| LVID;s (mm) | 2.902±0.09504 | 2.611±0.09318* |
| LVPW;d (mm) | 0.8222±0.02175 | 0.6703±0.05385* |
| LVPW;s (mm) | 1.229±0.06495 | 0.9139±0.05798 |
| **Morphometry** | n=13 | n=13 |
| BW (g) | 29.50±1.284 | 26.06±1.255 |
| VW/BW (mg/g) | 3.902±0.1393 | 4.184±0.1114 |
| AW/BW | 0.2011±0.009770 | 0.2078±0.02417 |
| Lu/BW | 4.902±0.2664 | 5.345±0.2230 |

Data are shown as mean ± SD. \*p<0.05, vs. WT at the same time point. Abbreviations: Echo= Echocardiography; BPM= beats per minute; EF= ejection fraction; IVS= interventricular septum; d= diastolic; s= systolic; LVID= left ventricular internal diameter; LVPW=left ventricular posterior wall thickness; BW= body weight; VW= ventricular weight; AW=atrial weight; Lu= lung weight.

**Table S3:** Echocardiography and morphometry four months after Tamoxifen administration.

|  | Male |  | Female |  |
| --- | --- | --- | --- | --- |
| | WT | αMHC-MerCreMer | WT | αMHC-MerCreMer |
| **Echo** | n=5 | n=4 | n=4 | n=4 |
| BPM | 450.6±27.28 | 446.3±25.86 | 471.3±6.290 | 451.3±5.406 |
| EF (%) | 57.68±3.073 | 60.62±3.461 | 40.27±2.845 | 48.10±4.927 |
| IVS; d (mm) | 0.86±0.049 | 0.849±0.053 | 0.8559±0.086 | 0.736±0.07 |
| IVS; s (mm) | 1.274±0.058 | 1.242±0.124 | 1.132±0.134 | 1.047±0.081 |
| LVID; d (mm) | 3.941±0.136 | 3.705±0.096 | 3.819±0.025 | 3.539±0.111* |
| LVID; s (mm) | 2.759±0.135 | 2.518±0.070 | 3.082±0.067 | 2.704±0.173 |
| LVPW; d (mm) | 0.805±0.013 | 0.6104±0.064* | 0.844±0.047 | 0.7302±0.084 |
| LVPW; s (mm) | 1.182±0.072 | 0.8941±0.074* | 1.289±0.120 | 0.934±0.1 |
| **Morphometry** | n=8 | n=5 | n=5 | n=8 |
| BW (g) | 32.38±1.114 | 29.14±1.389 | 24.90±0.871 | 24.13±1.524 |
| VW/BW (mg/g) | 3.905±0.186 | 3.897±0.234 | 4.304±0.059 | 4.129±0.179 |
| AW/BW | 0.204±0.012 | 0.1960±0.019 | 0.2617±0.052 | 0.1741±0.015 |
| Lu/BW | 4.366±0.202 | 4.778±0.185 | 5.762±0.374 | 5.699±0.282 |

Data are shown as mean ± SD. \*p<0.05 vs. WT at the same time point. Abbreviations: Echo= Echocardiography; BPM= beats per minute; EF= ejection fraction; IVS= interventricular septum; d= diastolic; s= systolic; LVID= left ventricular internal diameter; LVPW=left ventricular posterior wall thickness; BW= body weight; VW= ventricular weight; AW=atrial weight; Lu= lung weight.

**Table S4:** Echocardiography and morphometry one month after Tamoxifen administration.

|  | Male |  | Female |  |
| --- | --- | --- | --- | --- |
|  | Eef1a1/a2-WT | Eef1a1/a2-cKO | Eef1a1/a2-WT | Eef1a1/a2-cKO |
| **Echo** | n=6 | n=7 | n=4 | n=4 |
| BPM | 443.5±5.31 | 438.6±8.16 | 463±11.15 | 448.8±9.28 |
| EF (%) | 41.97±2.44 | 50.27±2.47* | 46.05±1.66 | 43.35±0.72 |
| IVS; d (mm) | 0.763±0.028 | 0.804±0.029 | 0.825±0.015 | 0.714±0.042 |
| IVS; s (mm) | 1.074±0.046 | 1.169±0.045 | 1.124±0.057 | 1.088±0.050 |
| LVID; d (mm) | 4.581±0.059 | 4.279±0.061** | 4.202±0.018 | 3.97±0.054* |
| LVID; s (mm) | 3.637±0.078 | 3.19±0.075** | 3.49±0.14 | 3.254±0.062 |
| LVPW; d (mm) | 0.874±0.031 | 0.786±0.030 | 0.737±0.049 | 0.689±0.047 |
| LVPW; s (mm) | 1.115±0.064 | 1.095±0.066 | 0.918±0.076 | 0.929±0.073 |
| **Morphometry** | n=3 | n=6 | n=3 | n=5 |
| BW (g) | 24.38±0.13 | 23.68±0.87 | 19.24±0.65 | 19.71±0.35 |
| VW/BW (mg/g) | 4.607±0.25 | 5.025±0.35 | 4.709±0.27 | 5.124±0.12 |
| AW/BW | 0.200±0.020 | 0.255±0.023 | 0.237±0.013 | 0.250±0.019 |
| Lu/BW | 5.804±0.11 | 7.199±0.41 | 6.846±0.56 | 6.644±0.44 |

Data are shown as mean ± SD. \*p<0.05, \*\*p<0.01 vs. WT at the same time point. Abbreviations: Echo= Echocardiography; BPM= beats per minute; EF= ejection fraction; IVS= interventricular septum; d= diastolic; s= systolic; LVID= left ventricular internal diameter; LVPW=left ventricular posterior wall thickness; BW= body weight; VW= ventricular weight; AW=atrial weight; Lu= lung weight.

**Table S5:** Echocardiography and morphometry four months after Tamoxifen administration.

|  | Eef1a1 -WT | Eef1a1-cKO |
| --- | --- | --- |
| **Echo** | n=10 | n=10 |
| BPM | 473.4±12 | 430.6±18 * |
| EF (%) | 50.42±2.513 | 47.08±2.431 |
| IVS;d (mm) | 0.880±0.038 | 0.825±0.043 |
| IVS;s (mm) | 1.209±0.056 | 1.14±0.042 |
| LVID;d (mm) | 4.085±0.12 | 3.885±0.13 |
| LVID;s (mm) | 3.163±0.12 | 2.988 ± 0.13 |
| LVPW;d (mm) | 0.754±0.022 | 0.742 ± 0.026 |
| LVPW;s (mm) | 1.039±0.041 | 0.960 ± 0.034 |
| **Morphometry** | n=10 | n=10 |
| BW (g) | 26.21±1.505 | 24.79±1.29 |
| VW/BW (mg/g) | 5.338±0.253 | 4.928±0.112 |
| AW/BW | 0.268±0.016 | 0.2736±0.02 |
| Lu/BW | 5.567±0.14 | 5.944±0.323 |

Data are shown as mean ± SD. \*p<0.05 vs. WT at the same time point. Abbreviations: Echo= Echocardiography; BPM= beats per minute; EF= ejection fraction; IVS= interventricular septum; d= diastolic; s= systolic; LVID= left ventricular internal diameter; LVPW=left ventricular posterior wall thickness; BW= body weight; VW= ventricular weight; AW=atrial weight; Lu= lung weight.

**Table S6:** Echocardiography and morphometry four months after Tamoxifen administration.

|  | Male |  | Female |  |
| --- | --- | --- | --- | --- |
|  | Eef1a1-WT | Eef1a1-cKO | Eef1a1-WT | Eef1a1-cKO |
| **Echo** | n=5 | n=5 | n=5 | n=5 |
| BPM | 465.6±16 | 442.6±36 | 492.2±18 | 422±11** |
| EF (%) | 54.25±3.538 | 43.73±2.251* | 46.58±2.92 | 49.65±2.47 |
| IVS; d (mm) | 0.887±0.041 | 0.854±0.048 | 0.864±0.094 | 0.805±0.068 |
| IVS; s (mm) | 1.261±0.071 | 1.135±0.053 | 1.083±0.061 | 1.144±0.064 |
| LVID; d (mm) | 4.137±0.16 | 4.134±0.23 | 3.961±0.077 | 3.707±0.13 |
| LVID; s (mm) | 3.211±0.16 | 3.257±0.21 | 3.048±0.079 | 2.795±0.13 |
| LVPW; d (mm) | 0.792±0.023 | 0.782±0.030 | 0.664±0.027 | 0.713±0.030 |
| LVPW; s (mm) | 1.098±0.042 | 0.986±0.032 | 0.898±0.063 | 0.942±0.055 |
| **Morphometry** | n=5 | n=5 | n=5 | n=5 |
| BW (g) | 24.38±0.13 | 23.68±0.87 | 30.07±0.867 | 28.48±2.11 |
| VW/BW (mg/g) | 4.607±0.25 | 5.025±0.35 | 5.895±0.335 | 5.073±0.158 |
| AW/BW | 0.200±0.020 | 0.255±0.023 | 0.300±0.012 | 0.296±0.030 |
| Lu/BW | 5.804±0.11 | 7.199±0.41 | 5.308±0.146 | 5.442±0.285 |

Data are shown as mean ± SD. \*p<0.05, \*\*p<0.01 vs. WT at the same time point. Abbreviations: Echo= Echocardiography; BPM= beats per minute; EF= ejection fraction; IVS= interventricular septum; d= diastolic; s= systolic; LVID= left ventricular internal diameter; LVPW=left ventricular posterior wall thickness; BW= body weight; VW= ventricular weight; AW=atrial weight; Lu= lung weight.

**Table S7:**
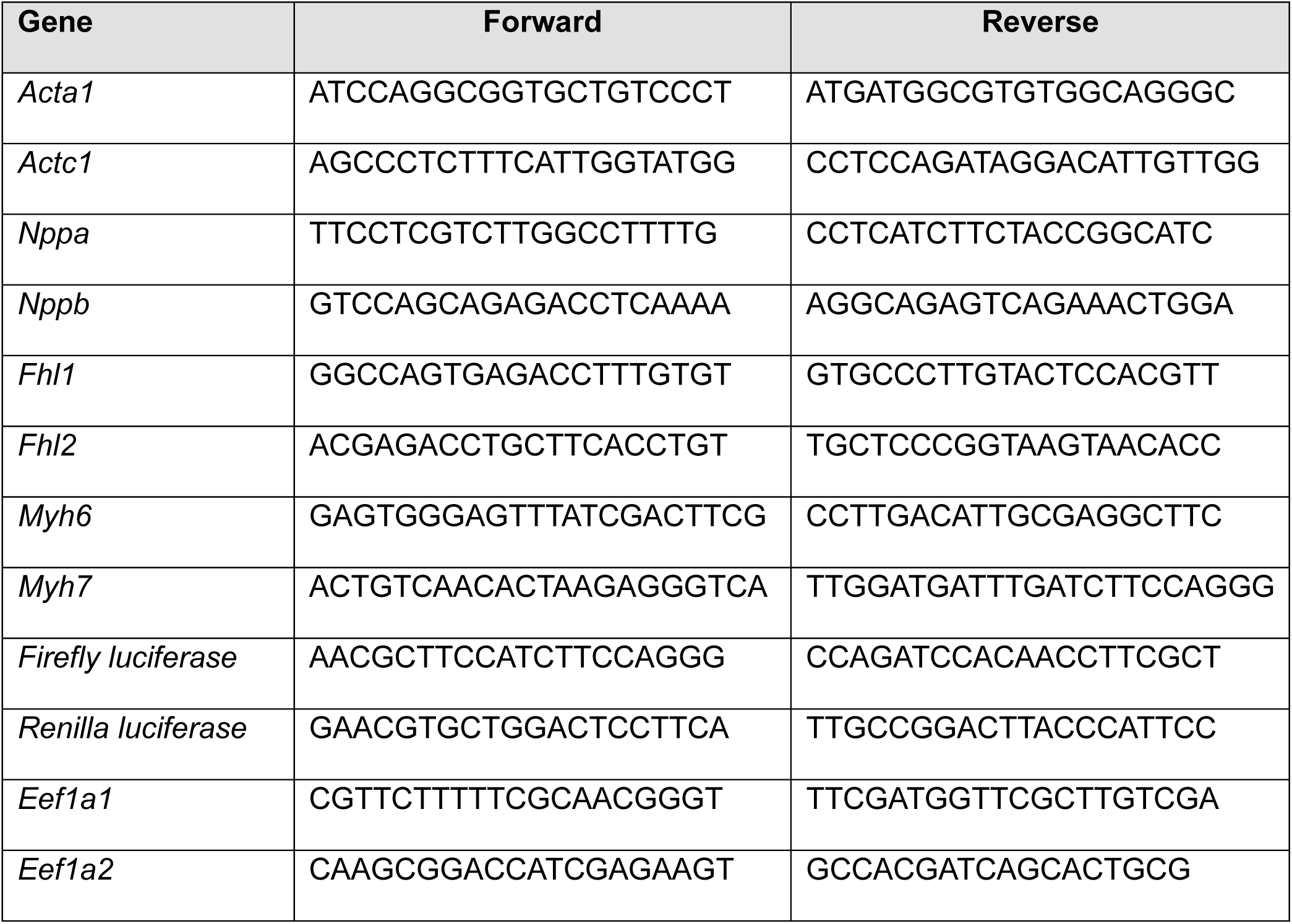
List of primers used for qRT-PCR (5’-3’).

**Table S8:** List of antibodies used for Western blotting (WB) and immunofluorescence (IF).

| **Antibody** | **Vendor** | **Catalog number** | **Dilution** |
| --- | --- | --- | --- |
| eEF1A1 | Proteintech | 11402-1-AP | WB: 1:2000 |
| eEF1A2 | Proteintech | 16091-1-AP | WB: 1:2000 |
| RpS6 (5G10) | Cell Signaling | # 2217 | WB: 1:1000, IF: 1:100 |
| pRpS6 (Ser235/236) (D57.2.2E) | Cell Signaling | # 4858 | WB: 1:1000, IF: 1:100 |
| eEF2 | Cell Signaling | #2332 | Cell Signaling |
| peEF2 (Thr56) | Cell Signaling | #2331 | WB: 1:1000 |
| p62/SQSTM1 | Sigma | P0067 | WB: 1:2000, IF: 1:200 |
| LC3 | Proteintech | 14600-1-AP | WB: 1:1000 |
| eIF2α | Cell Signaling | #9722 | WB: 1:1000 |
| peIF2α (Ser51) (D9G8) | Cell Signaling | #3398 | WB: 1:1000 |
| Atg7 | Cell Signaling | #2631 | WB: 1:1000 |
| Beclin-1 (D40C5) | Cell Signaling | #3495 | WB: 1:1000 |
| EDF1 | Proteintech | 12419-1-AP | WB: 1:2000 |
| α-Actinin (EA-53) | Sigma | A7811 | IF: 1:200 |
| Puromycin Antibody (12D10) | Sigma | MABE343 | IF: 1:200 |
| Wheat Germ Agglutinin (WGA) | Invitrogen | W21405 | IF: 1:100 |
| GAPDH (14C10) | Cell Signaling | #2118 | WB: 1:1000 |
| Ubiquitinated proteins (FK2) | Sigma | 04-263 | IF: 1:100 |
| Anti-mouse IgG, HRP-linked | Cell Signaling | #7076 | WB: 1:1000 |
| Anti-rabbit IgG, HRP-linked | Cell Signaling | #7074 | WB: 1:1000 |
| Anti-rabbit IgG (H+L), F(ab')<sub>2</sub> Fragment (Alexa Fluor® 488 Conjugate) | Cell Signaling | #4412 | IF: 1:200 |
| Anti-rabbit IgG (H+L), F(ab')<sub>2</sub> Fragment (Alexa Fluor® 555 Conjugate) | Cell Signaling | #4413 | IF: 1:200 |
| Anti-mouse IgG (H+L), F(ab')<sub>2</sub> Fragment (Alexa Fluor® 488 Conjugate) | Cell Signaling | #4408 | IF: 1:200 |
| Anti-mouse IgG (H+L), F(ab')<sub>2</sub> Fragment (Alexa Fluor® 555 Conjugate) | Cell Signaling | #4409 | IF: 1:200 |

**Figure S1.**
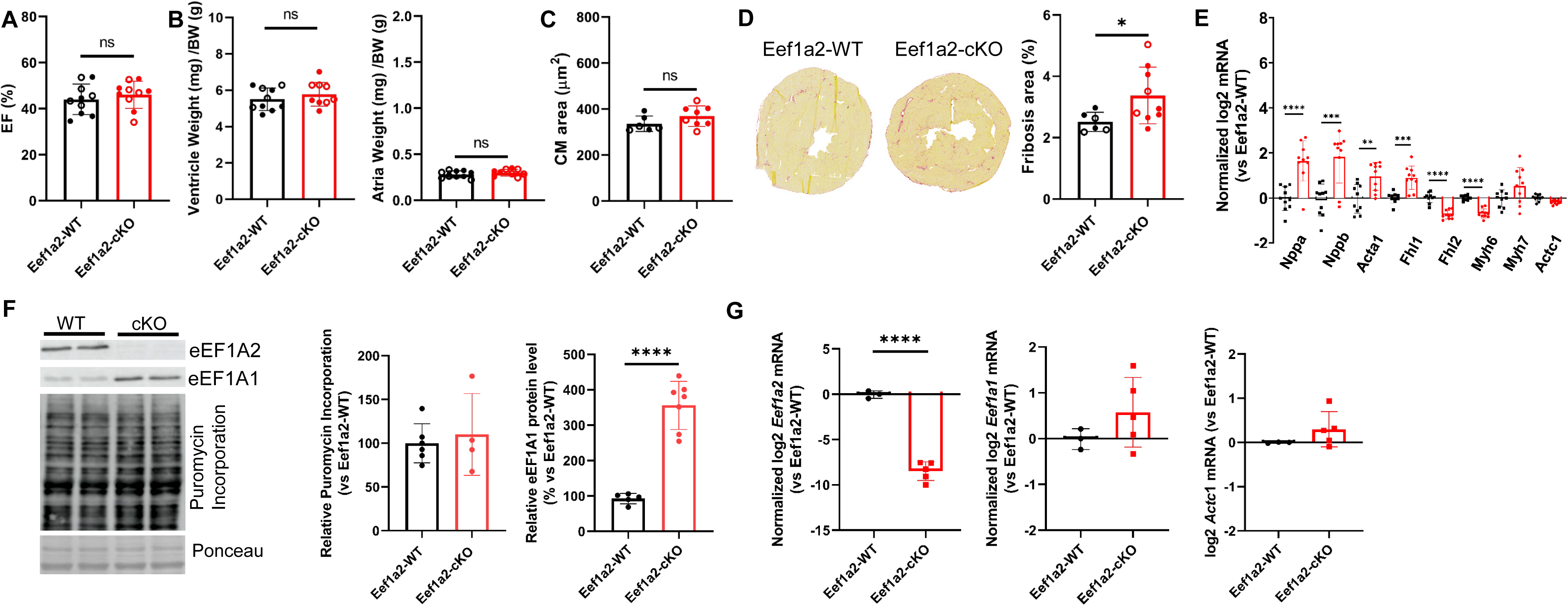
Effects of cardiomyocyte-specific Eef1a2 deletion at 1 month post gene-deletion. (**A**) Left ventricular ejection fraction of Eef1a2-WT (n = 10) and Eef1a2-cKO (n = 10) mice.(**B**) Ventricular (left) and atrial (right) weight normalized to body weight in Eef1a2-WT (n = 10) and Eef1a2-cKO (n = 9) mice at 1 month after Eef1a2 deletion. (**C**) Quantification of cardiomyocyte cross sectional area in cardiac sections stained with wheat germ agglutinin (WGA) from Eef1a2-WT (n = 10) and Eef1a2-cKO (n = 10) mice.(**D**) Representative Sirius Red staining of cardiac fibrosis and corresponding quantification of fibrotic area from Eef1a2-WT (n = 6) and Eef1a2-cKO (n = 9) mice. *P < 0.05, unpaired two-tailed t-test. (**E)** Normalized mRNA expression levels of selected genes in cardiac tissue from Eef1a2-WT (n = 10) and Eef1a2-cKO (n = 10) mice. **P < 0.01, ***P < 0.001, ****P < 0.0001; unpaired two-tailed t-test.(**F**) Representative Western blot analysis of isolated cardiomyocytes and quantification of puromycin incorporation and eEF1A1 protein levels in isolated cardiomyocytes from Eef1a2-WT (n = 5) and Eef1a2-cKO (n = 4 for puromycin incorporation and n=7 for Eef1a1) mice from independent isolations. ****P < 0.0001; unpaired two-tailed t-test. (**G**) mRNA expression levels of *Eef1a2, Eef1a1* and *Actc1* in 5 independent cardiomyocyte isolations from Eef1a2-cKO mice. *Actc1* mRNA expression is used as control. ****P < 0.0001; unpaired two-tailed t-test. Filled circles denote males; open circles denote females.

**Figure S2.**
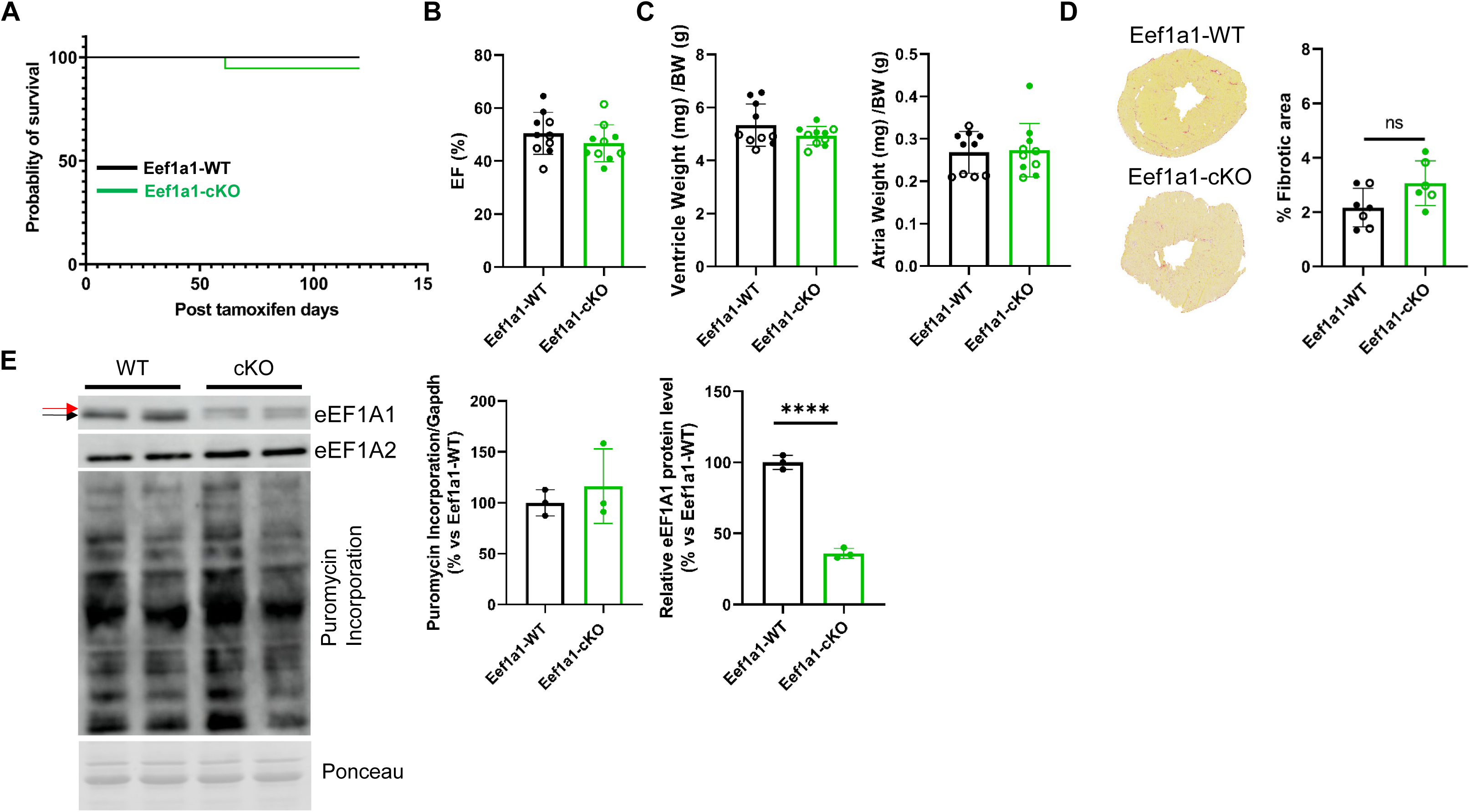
Effects of cardiomyocyte-specific Eef1a1 deletion. (**A**) Kaplan–Meier survival curves of Eef1a1-WT (n = 10, 10/10 survived) and Eef1a1-cKO (n = 15, 14/15 survived) mice. (**B**) Left ventricular ejection fraction (EF) of Eef1a1-WT and Eef1a1-cKO. (**C**) Ventricular (left) and atrial (right) weight normalized to body weight in Eef1a1-WT and Eef1a1-cKO mice. (**D**) Representative Sirius Red staining of cardiac fibrosis and corresponding quantification of fibrotic area from Eef1a1-WT (n=7) and Eef1a1-cKO (n=6) mice. (**E**) Representative Western blot analysis of isolated cardiomyocytes at 1 months after Eef1a1 deletion and quantification of puromycin incorporation, eEF1A1 and eEF1A2 protein level in isolated cardiomyocytes from Eef1a1-WT (n = 3) and Eef1a1-cKO (n = 3) mice from independent isolations at 1 month after Eef1a1 deletion. The black arrow denotes eEF1A1, whereas then red arrow denotes eEF1A2. ****P < 0.0001; unpaired two-tailed t-test. Data in B-D were obtained 4 months after gene deletion. Filled circles indicate males, and open circles indicate females.

**Figure S3.**
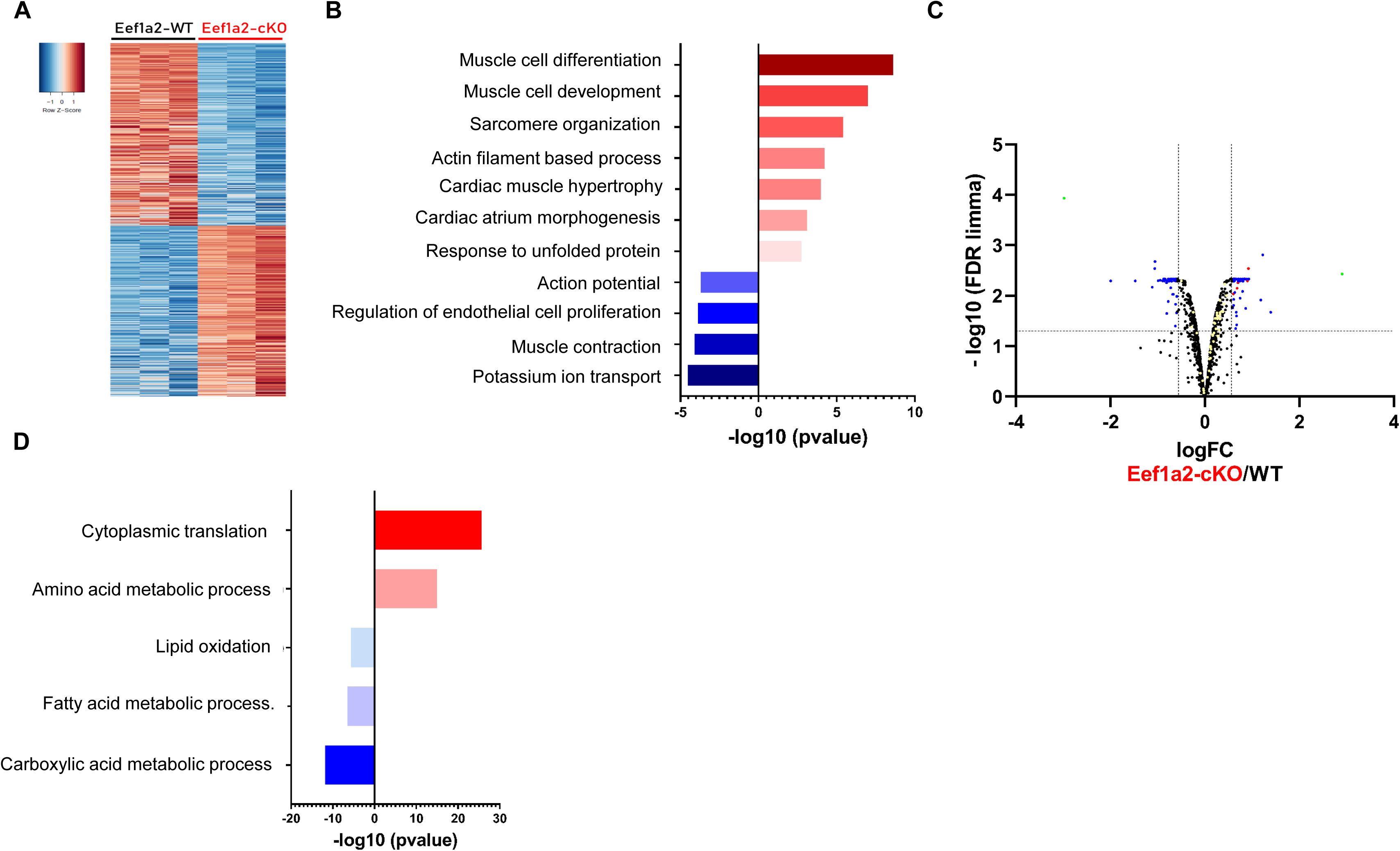
Effects of 1 month of Eef1a2 deletion on mRNA and protein expression in cardiac tissue. (**A**) Heat map representation of differentially expressed genes from RNA sequencing. (**B**) Gene Ontology (GO) analysis of regulated genes identified in (A). Upregulated GO terms in Eef1a2-cKO are shown in red and downregulated GO terms in blue. (**C**) Volcano plot of proteomics results showing fold-change (FC; log₂) and P value of protein abundance changes in cardiac tissue from WT (n = 3) and Eef1a2-cKO (n = 3) mice at 1 month after Eef1a2 deletion. Significantly regulated proteins are highlighted in blue, while significant ribosomal proteins or elongation factors are highlighted in red; yellow labels unsignificant ribosomal proteins or elongation factors. (**D**) Gene Ontology (GO) analysis of significant proteins identified in (C), shown as −log₁₀(P value). Upregulated GO terms in Eef1a2-cKO are shown in red and downregulated GO terms in blue.

**Figure S4.**
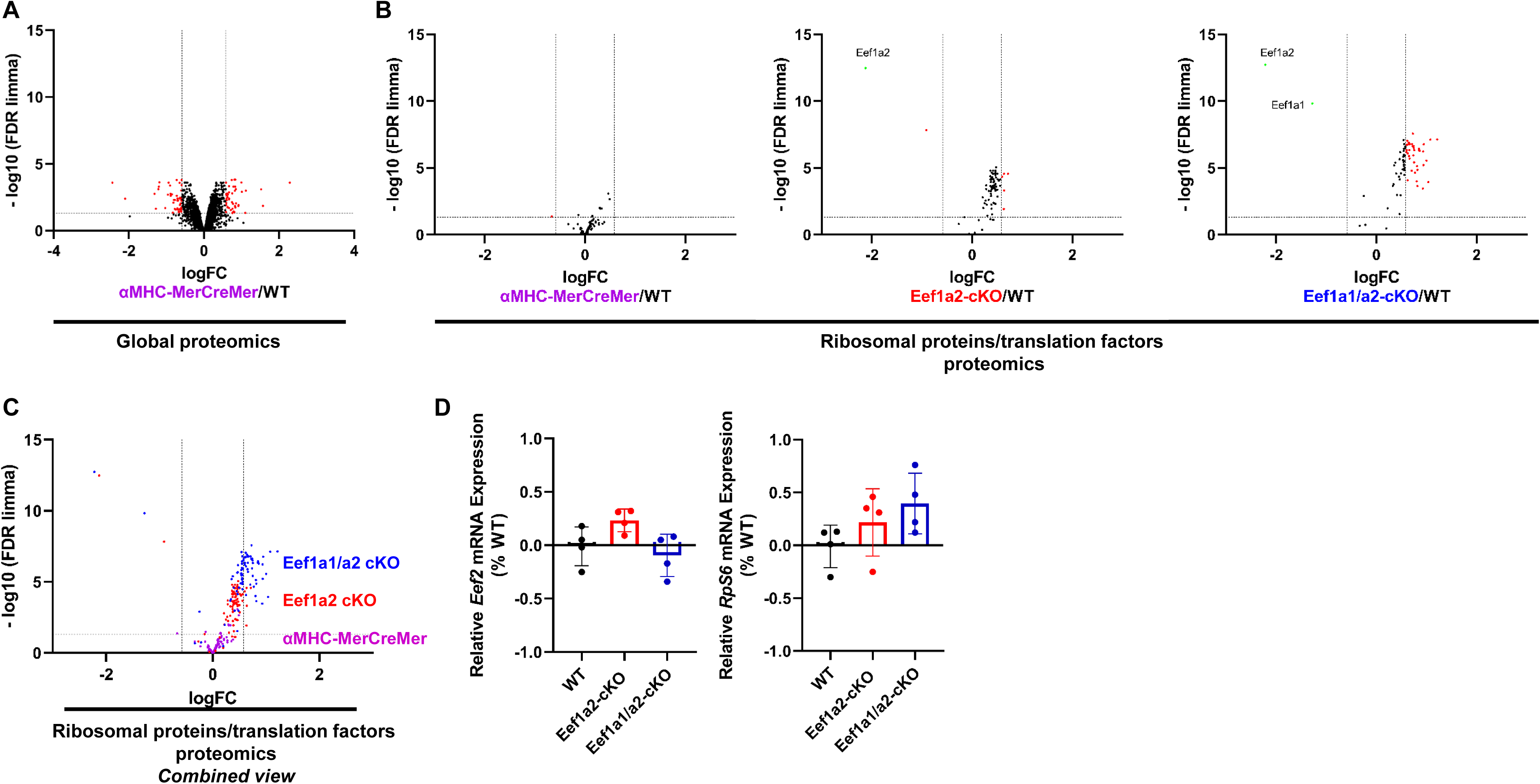
Effects of 1 month of Eef1a2 and Eef1a1 deletion on ribosomal and translation factor protein and mRNA expression in isolated cardiomyocytes. (**A**) Volcano plot showing fold-change (FC; log₂) and P value (−log₁₀) of protein abundance changes in isolated cardiomyocytes from WT (n = 4) and MerCreMer (n = 4) mice at 1 month after tamoxifen injection. (**B**) Volcano plot showing fold-change (FC; log₂) and P value (−log₁₀) of ribosomal and translation factor changes in in isolated cardiomyocytes from WT (n = 4), αMHC-MerCreMer (n=4), Eef1a2-cKO (n=4) and Eef1a1/a2-cKO (n = 4) mice at 1 month after tamoxifen injection. Significantly regulated proteins are highlighted in red. (**C**) Volcano plot representation of the combined data shown in (B). (**D**) mRNA expression levels of Eef1a1 and Rps6 in 4 independent cardiomyocyte isolations in the indicated mice.

**Figure S5.**
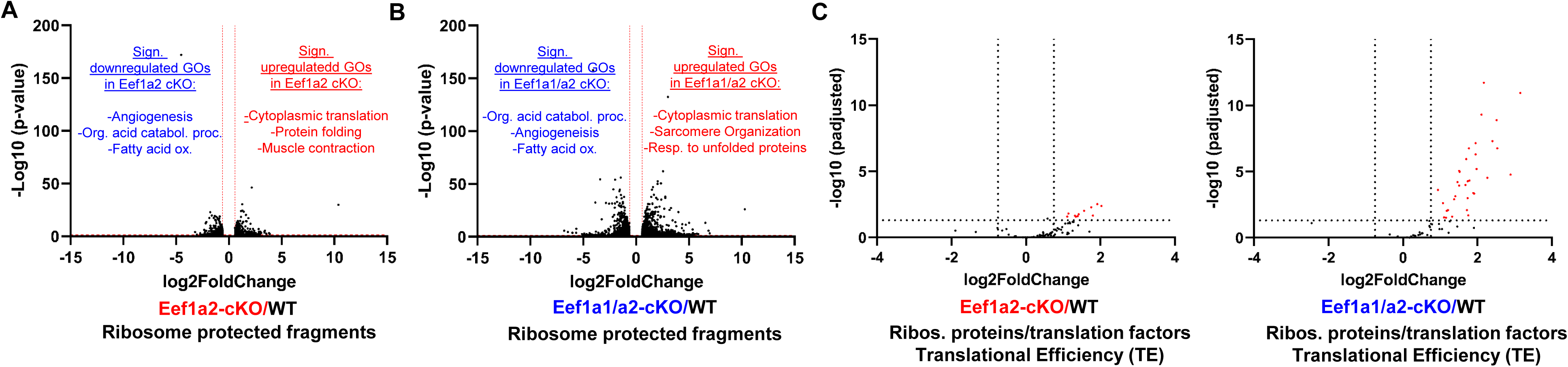
Effects of Eef1a on translation in cardiac tissue. (**A**) Volcano plot showing fold-change (FC; log₂) and P value (−log₁₀) of changes in ribosome protected fragments in cardiac tissue from WT (n = 4) and Eef1a2-cKO (n = 4) mice at 1 month after Eef1a2 deletion. Significantly upregulated GO terms in Eef1a2-cKO are shown in red, whereas downregulated terms are shown in blue. (**B**) Volcano plot showing fold-change (FC; log₂) and P value (−log₁₀) of changes in ribosome protected fragments in cardiac tissue from WT (n = 4) and Eef1a1/a2-cKO (n = 4) mice at 1 month after Eef1a1/a2 deletion. Significantly upregulated GO terms in Eef1a1/a2-cKO are shown in red, whereas downregulated terms are shown in blue. (**C**) Volcano plots showing fold change (FC; log₂) and P value (−log₁₀) of changes in translation efficiency of ribosomal proteins and translation factors in cardiac tissue from WT (n = 4), Eef1a2-cKO (n = 4), and Eef1a1/a2-cKO (n = 4) mice at 1 month after tamoxifen injection. Regulated genes are highlighted in red.

**Figure S6.**
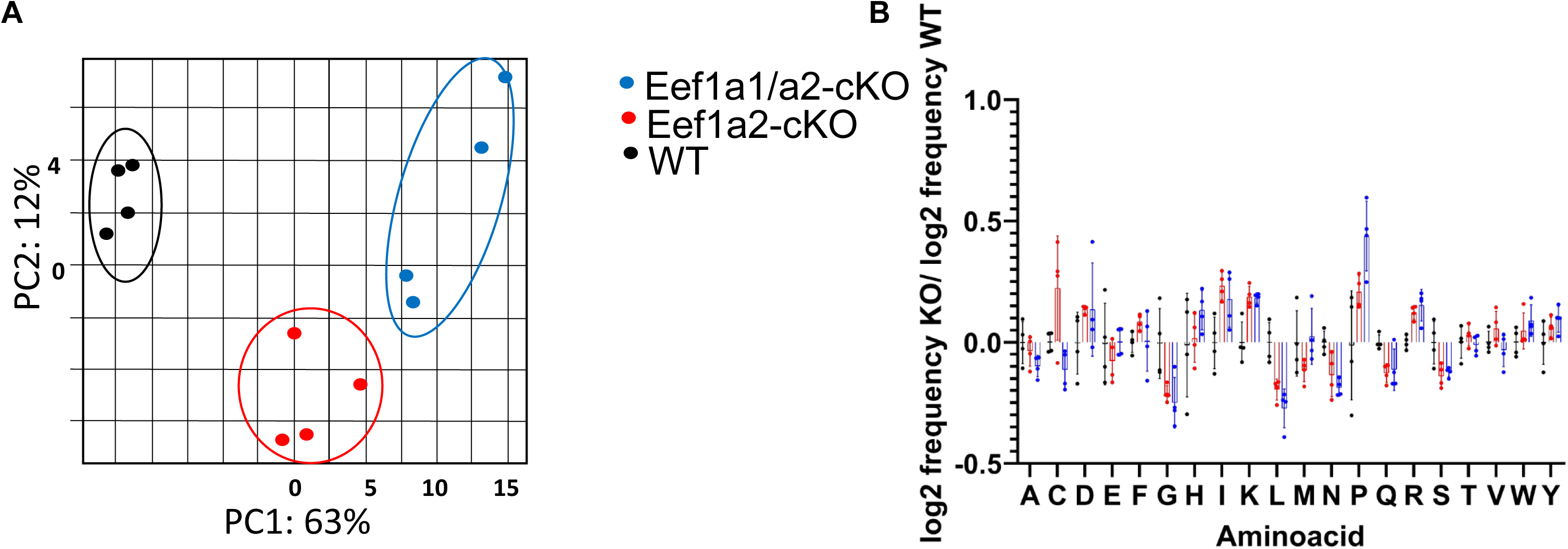
Codon occupancy in cardiac tissue 1 month after Eef1a2 or Eef1a1/a2 deletion. (**A**) Principal Component Analysis (PCA) showing clustering of the different experimental groups.(**B**) P-site amino acid occupancy of individual amino acids in cardiac tissue.

**Figure S7.**
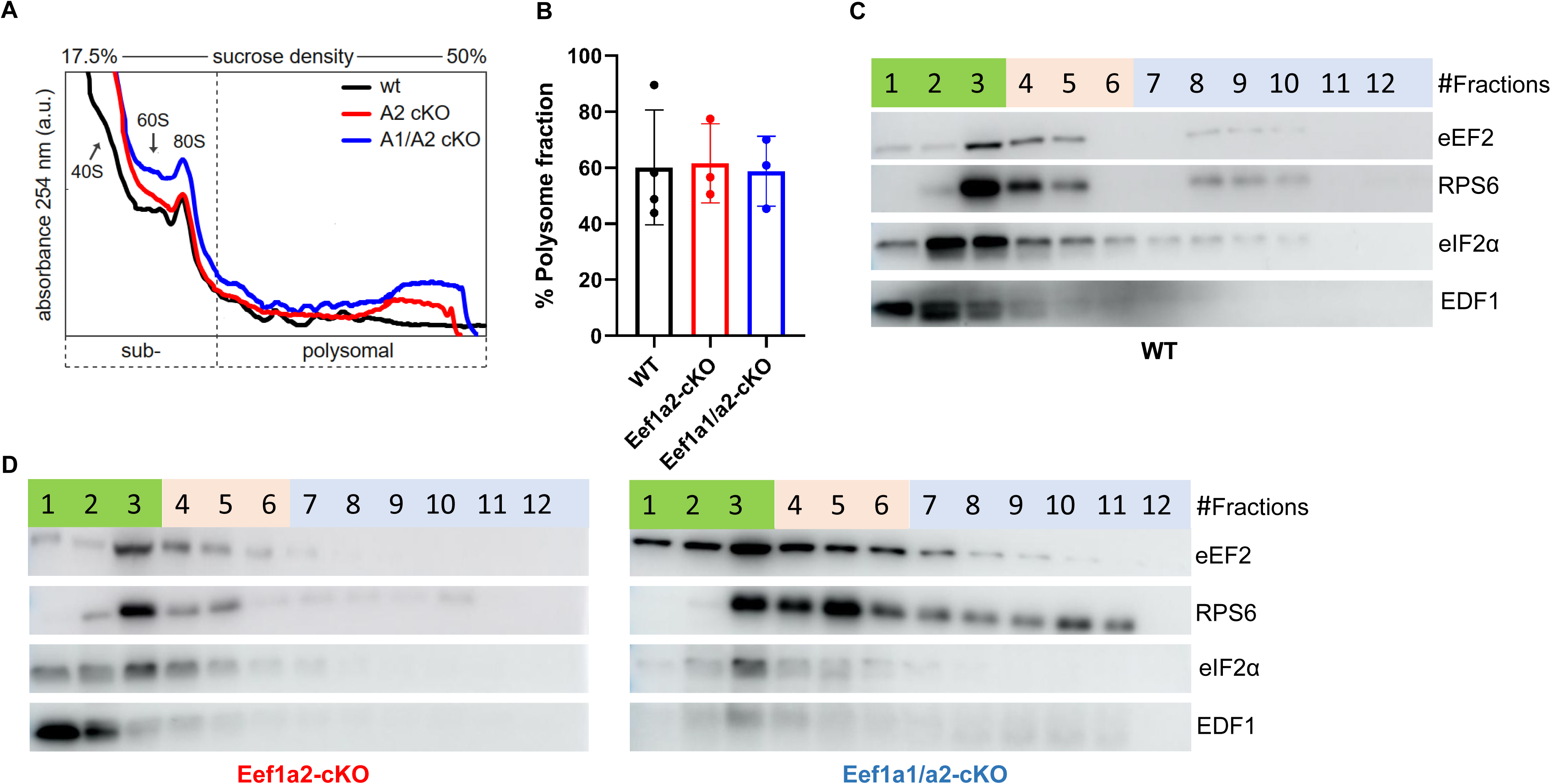
Polysome profiling of WT, Eef1a2-cKO, and Eef1a1/a2-cKO from cardiac tissue 1 month after tamoxifen. (**A**) Representative monosome/polysome profiles.(**B**) Quantification of polysome fractions from WT (n = 3), Eef1a2-cKO (n = 3), and Eef1a1/a2-cKO (n = 3) cardiac tissue.(**C**) Representative Western blot of eEF2, RPS6, eIF2α, and EDF1 in fractions collected from WT cardiac tissue. (**D**) Same analysis as in (C) from Eef1a2-cKO (left) and Eef1a1/a2-cKO (right) cardiac tissue.

**Figure S8.**
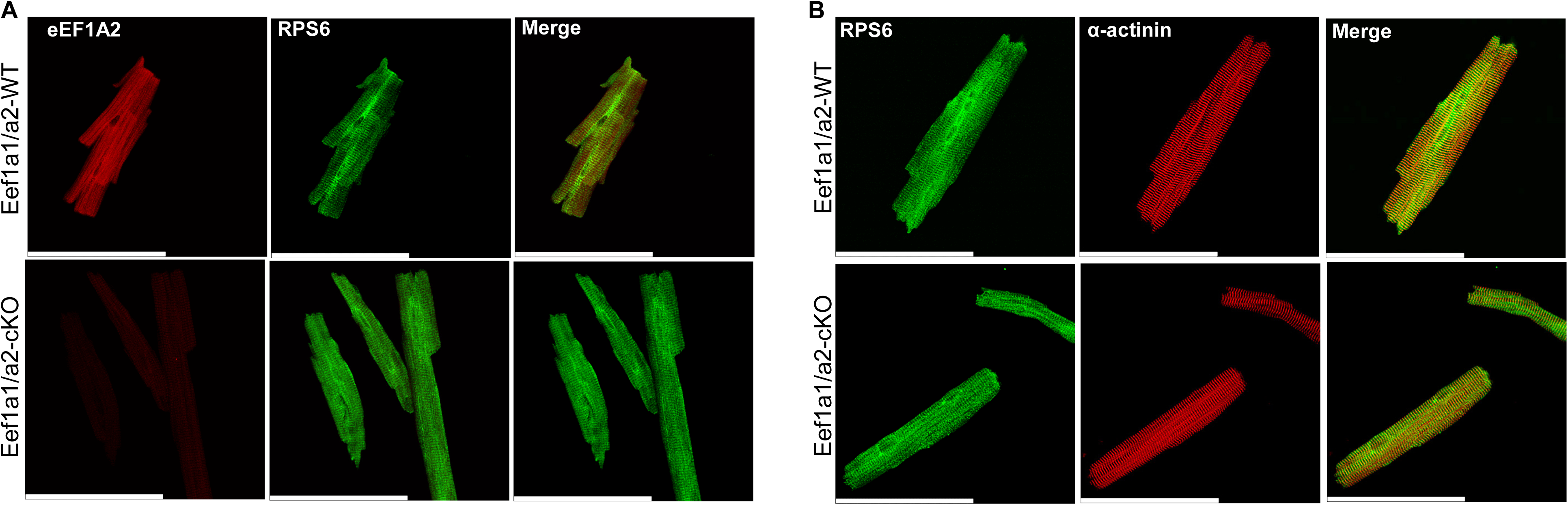
Representative co-immunostaining in isolated cardiomyocytes from Eef1a1/a2-WT or Eef1a1/a2-cKO mice at 1 month after Tamoxifen. **(A)** Staining for eEF1A2 and RPS6 in isolated cardiomyocytes from the indicated mice. **(B)** Staining for RPS6 and α-actinin in isolated cardiomyocytes from the indicated mice. Scale bars: 100µm.

**Figure S9.**
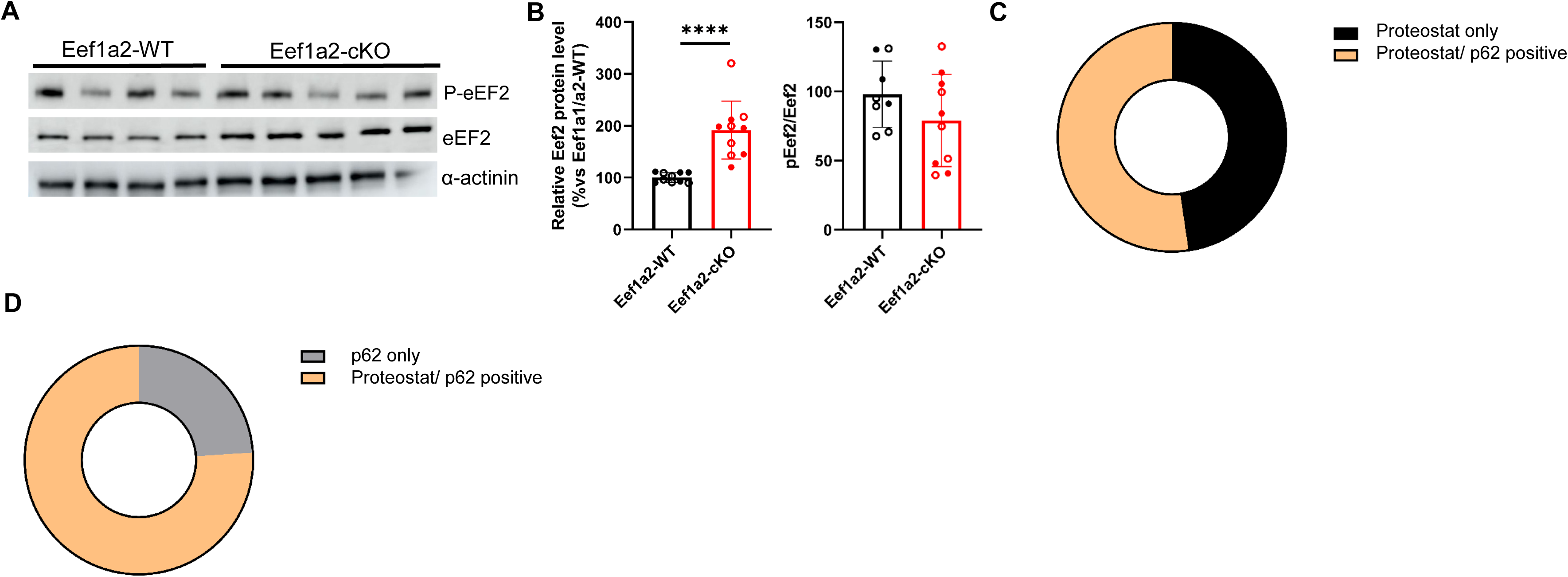
Western blot and autophagy analysis in Eef1a2-cKO cardiac tissue. (**A**) Representative Western blot analysis of the indicated proteins in cardiac tissue from Eef1a2-WT and Eef1a2-cKO mice at 2 months after Eef1a2 deletion. (**B**) Quantification of protein levels shown in (A) from Eef1a2-WT (n = 9) and Eef1a2-cKO (n = 10) mice. Filled circles denote males; open circles denote females. ****P < 0.0001; unpaired t-test. (**C**) Circle representation of Proteostat-positive cardiomyocytes alone or co-stained with p62 in Eef1a2-cKO hearts. (**D**) Circle representation of p62-positive cardiomyocytes alone or co-stained with Proteostat in Eef1a2-cKO hearts.

**Figure S10.**
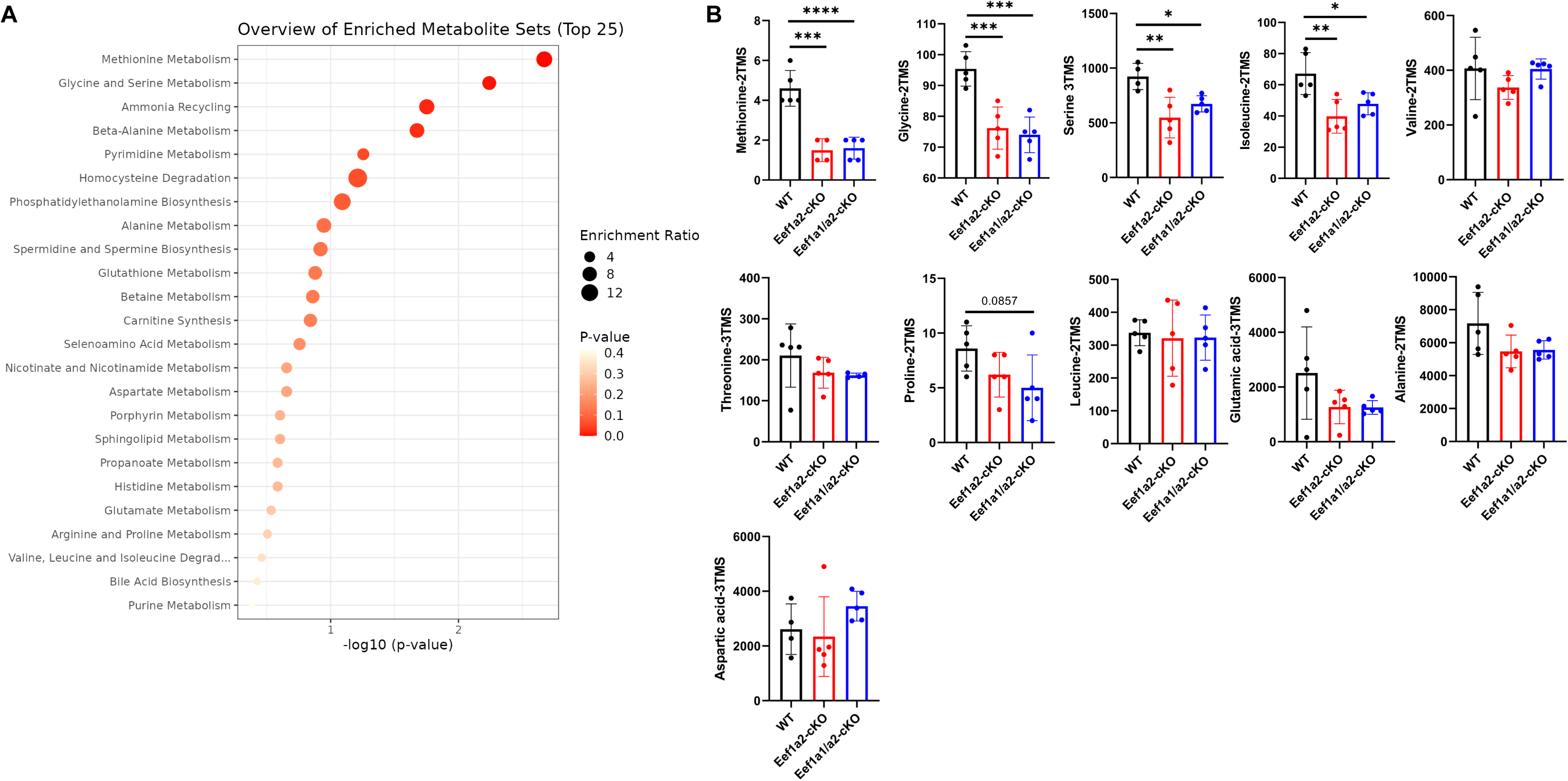
Metabolomics in heart tissue from wild-type (WT), Eef1a2-cKO or Eef1a1/a2-cKO mice at 1 month after Tamoxifen. **(A)** Overview of significantly downregulated metabolite sets in Eef1a2-cKO versus WT mice. No significant upregulated metabolites were detected. **(B)** Quantification of detected amino acids in the myocardium of indicated mice. N=5/genotype. *p<0.05; **p<0.01; ***p<0.001; ****p<0.0001

**Figure S11.**
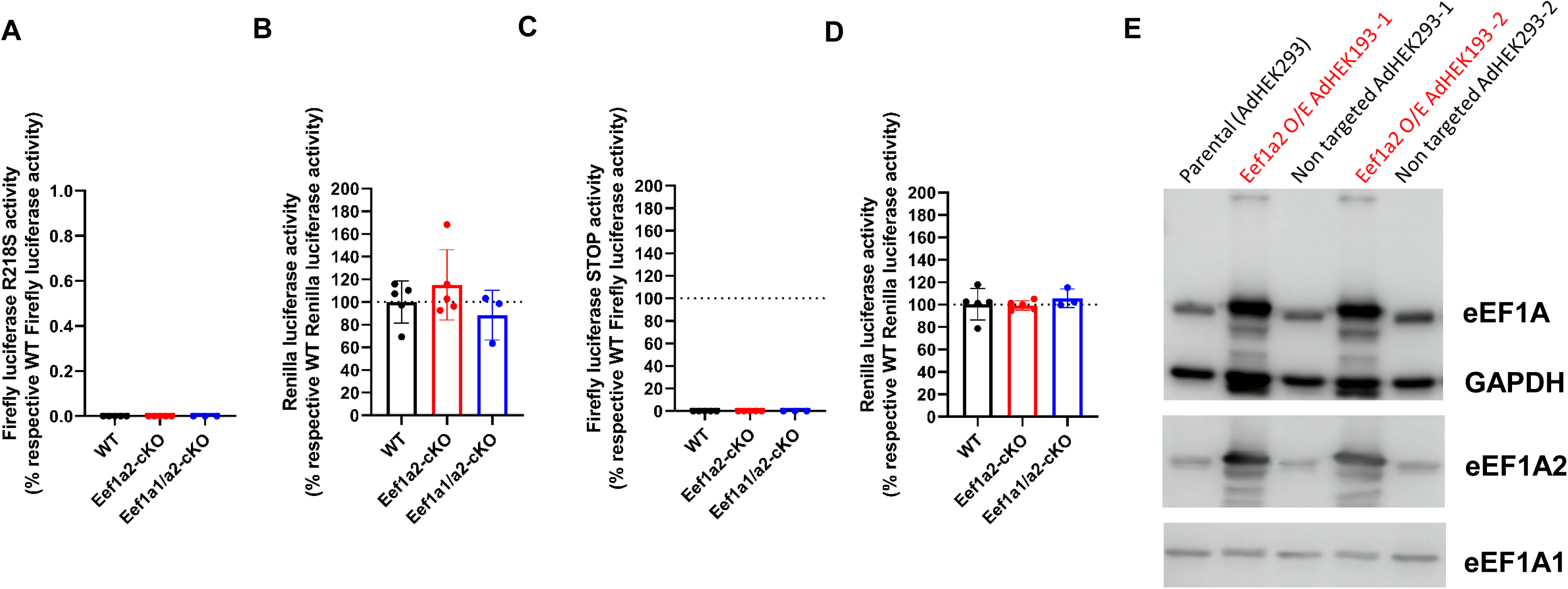
Effect of eEF1A2 on protein translation fidelity. **(A**) Relative firefly R218S luciferase activity measured in isolated cardiomyocytes from Eef1a2-cKO (n = 5) or Eef1a1/a2-cKO (n = 5) mice, normalized to their respective wild-type controls.(**B**) Relative Renilla luciferase activity from the samples in (A).(**C**) Relative firefly STOP luciferase activity measured in isolated cardiomyocytes from Eef1a2-cKO (n = 5) or Eef1a1/a2-cKO (n = 5) mice, normalized to their respective wild-type controls.(**D**) Relative Renilla luciferase activity from the samples in (C).(**E**) Representative Western blot of Eef1a2-overexpressing and non-targeted clones used in the folding/unfolding assay.

**Figure S12.**
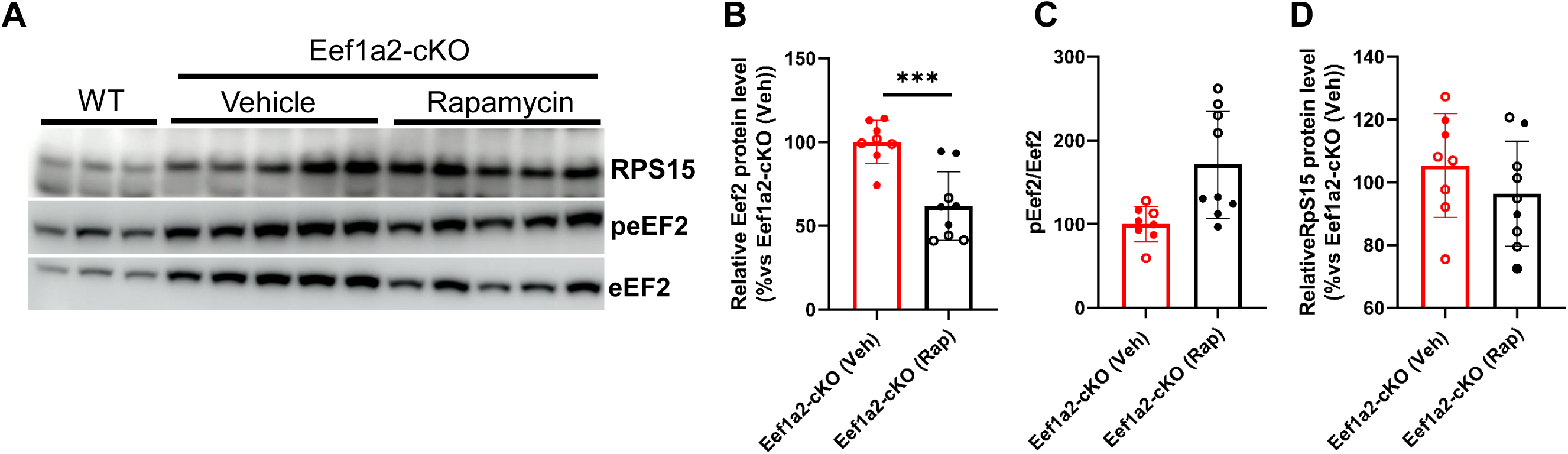
Western blot analysis in cardiac tissue from vehicle- or Rapamycin-treated Eef1a2-cKO mice. **(A**) Representative Western blot analysis of RPS15, eEF2, and peEF2 in cardiac tissue from vehicle- or Rapamycin-treated Eef1a2-cKO mice. (**B, C and D**) Quantification of protein expression shown in (A) in vehicle-treated (n = 8) or Rapamycin-treated (n = 9) Eef1a2-cKO mice. ***P < 0.001, unpaired two-tailed t-test.

